# Mathematical Model for the Progression of Rhegmatogenous Retinal Detachment (RRD)

**DOI:** 10.1101/2025.08.03.668360

**Authors:** William Ebo Annan, Emmanuel Asante-Asamani, Diana White

## Abstract

The separation of the neural layer (NL) and the retinal pigmented epithelium (RPE), referred to as retinal detachment (RD), is a disease of vertebrate eyes affecting nearly twenty eight thousand individuals in the United States annually. The rate at which RD progresses—especially in response to constant eye movement—and the factors influencing this progression remain poorly understood. This lack of quantitative insight contributes to delays in treatment which can lead to permanent vision loss. In this work, we develop a mathematical model to investigate the progression of retinal detachment over two saccadic eye rotation. We explore how various model parameters—describing fluid properties, biomechanical properties of the retina, molecular properties of the bond between the NL and the RPE, as well as geometric and physical properties of the eye—affect disease progression.

## 1. Introduction

The retina is the innermost tissue layer of the vertebrate eye, consisting of two main layers: the neural layer (NL) and the retinal pigmented epithelium (RPE)[55]. Retinal detachment (RD) refers to the separation of these two layers. In general, RD occurs when fluid accumulates beneath the NL, thereby causing separation. There are three types of RD: exudative (ERD), tractional (TRD), and rhegmatogenous retinal detachment (RRD), with RRD being the most prevalent [71]. RRD typically arises from a tear or break in the retina, allowing fluid from the vitreous chamber to enter the subretinal space (the space between the NL and the RPE). The accumulation of fluid in this space primarily drives the separation of the NL from the RPE. During saccadic eye movements (*rapid shifts of the eyes from one point to another*), the vitreous humor moves, exerting pressure on the detached NL and potentially exacerbating the detachment.

Various factors can precipitate RD, including lattice degeneration— a condition where the peripheral retina becomes thin and more susceptible to breaks [13]; advancing age; intense physical activity [43]; the shape of the eyeball (*myopia*) [21, 48]; ocular trauma [12]; inflammatory diseases such as diabetes [65, 25]; genetic predisposition [48]; and a history of posterior vitreous detachment (PVD) [26, 32, 6, 46].

Several studies have observed that photoreceptor cells gradually lose their normal function as detachment progresses [35, 73, 10, 15]. The outer segment (OS) of photoreceptor cells (*rods and cones*) comprises stacks of membranous structures called discs embedded with photosensitive proteins essential for converting light energy into electrical signals perceived by the brain to enable vision [38, 63, 23, 30]. Upon detachment, photoreceptor cells are deprived of nutrients and essential metabolic support, leading to shortening of the outer segment, impaired cellular function, cell death, and eventual permanent vision loss [41, 58, 7]. In the United States, approximately 10 to 18 individuals per 100,000 population develop RRD annually [69, 59], with older adults at greater risk. Age has been shown to correlate with an increased likelihood of retinal detachment [43, 21, 27], and both retinal thickness [39] and elasticity [45, 14] have been observed to decrease with age. However, to the best of our knowledge, no prior studies have investigated how these changes influence the progression of detachment.

Current ophthalmological examinations can detect the presence of RD, especially when retinal breaks or holes are involved, but they cannot reliably estimate the duration of the detachment or how rapidly it progresses [34]. As a result, patients with RD might be scheduled for treatment after significant photoreceptor damage has already occurred. In such cases, even if the NL is successfully reattached to the RPE, the photoreceptor cells may not regain their normal function, leading to permanent vision loss [1, 29].

In this study, we develop a retinal progression–fluid structure interaction (RP-FSI) model to track the progression of RRD. The model comprises a set of equations describing fluid flow, the motion of the retina in response to the fluid, and the distribution of adhesion bonds between the NL and the RPE that restrain retinal motion. The fluid and the structure components of the model are coupled through a Dirac delta function.

The concept of Fluid Structure Interaction (FSI), also known as the immersed boundary method, was first introduced by Charles Peskin [53] in 1972 to simulate the interaction between blood flow and the flexible heart structure, specifically modeling fluid motion within the heart and its coupling with elastic tissue deformations. Since then, different authors have used FSI models to study interactions between fluid and deformable structures in many application contexts [68, 44, 54, 74, 62, 36]. In 2018, Natali et al. [52] applied an FSI model to study retinal detachment, specifically looking at the propensity for further detachment at the clamped edge. However, their modeling framework did not account for the adhesion bonds between the NL and the RPE, as the focus of that study was on assessing the likelihood of further detachment rather than quantifying actual detachment. To the best of our knowledge, no other mathematical model has been developed to examine the progression of RD. This work therefore aims to compute the actual detachment that can occur within a cycle or a complete eye rotation (defined to be the movement of the eyeball from the forward looking direction to either right or left and returning to the initial position). To capture the detachment in this time frame during eyeball rotation, necessitates incorporating the mechanics of adhesion bond rupture between the NL and the RPE.

Existing fluid solvers were mostly developed *ad hoc* for specific problems, making it difficult to integrate the adhesion model into the existing FSI packages. We therefore developed a computational algorithm in MATLAB to solve the model: the adhesion bond model is solved using an implicit upwind scheme, the retinal motion using a finite difference approximation, and the fluid flow using the *P*_2_ projection method introduced by Chorin in 1968 [18], and later adapted by Kim and Moin for finite volume formulations on staggered grids [42].

Our goal is to investigate how the progression of RRD is influenced by several factors, including fluid properties, biomechanical properties of the retina, and physical and geometric characteristics such as the average rate of eye rotation, the elevation of the detached retina, and the initial detached length. In our analysis, we focused on one cycle or complete eye rotation and examined how the stated factors impact the progression of RD. We quantify the extent of detachment that can occur during this complete eye rotation as various model parameters are varied. We observed that increasing the speed of eye rotation, the initial detached length, the elevation of the detached retina, and the unbinding affinity of the adhesion proteins resulted in greater detachment, while increasing retinal thickness, retinal elasticity, and the strength of adhesion bonds between the NL and the RPE reduced the progression of detachment.

The rest of this work is organized as follows. In Section 2, we present the dimensional form of the RP-FSI model, state the initial and boundary conditions, and then derive its nondimensionalized form. This nondimensional form reduces the number of parameters and facilitates numerical implementation. In Section 3, we provide a brief overview of the numerical algorithm used to simulate the RP-FSI model (Detailed implementation strategy and discretization can be found in the supplementary material). We also present a table of the base parameter values and explain the rationale behind their selection. Section 4 presents the results and discussion. We begin with a convergence analysis of the numerical method, followed by results showing the flow field and detachment profile using the base parameter values. Finally, we examine the influence of individual model parameters on the progression of detachment by varying one parameter at a time while keeping the others fixed. The conclusions of this work are summarized in Section 5.

## 2. Governing Equations

When a tear or hole forms in the retina, fluid from the vitreous passes through and accumulates between the neural layer (NL) and the retinal pigmented epithelium (RPE), thereby causing separation between the two layers—a condition known as rhegmatogenous retinal detachment (RRD). Constant rotation of the eyeball sets the vitreous fluid in motion, exerting pressure on the retina and causing further detachment. To study this interaction, we develop a retinal progression–fluid-structure interaction (RP-FSI) model to investigate the progression of RRD, which results from interactions between the vitreous fluid and the detached retina. This model builds upon established fluid-structure interaction (FSI) frameworks and also incorporates the binding and unbinding dynamics of adhesion proteins between the NL and the RPE to capture how the adhesion bonds are broken, resulting in further detachment. The RP-FSI modeling framework is similar to the FSI modeling approach used by several authors to investigate the interaction between fluid and immersed structures [52, 44, 54, 62, 36].

### 2.1. Fluid and Retina Model

In typical FSI models, the fluid domain is described in Eulerian coordinates, denoted by **x** ∈ ℝ^2^, while the material or structural domain (here, the retina) is described in Lagrangian coordinates, denoted by **X** : [*a, b*] × [0, *T*] →ℝ^2^. Figure 1A shows a two-dimensional surface view of the retina with a hole and a small detached region around it, while Figure 1B shows a cross-sectional view representing the computational domain used in the study. In the RP-FSI model, the motion of the detached retina is influenced by adhesive interactions between the NL and the RPE.

**Figure 1:**
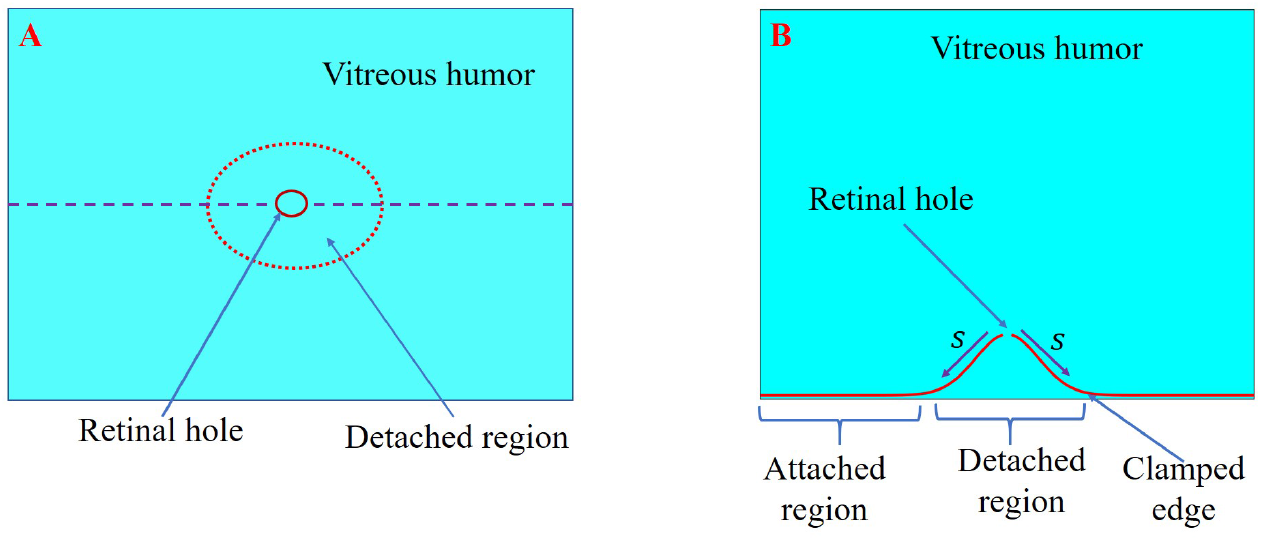
Computational domain: (A) Two-dimensional surface view of the retina showing a developed hole and a small detached region around it. (B) Cross-sectional view of the detached retina representing the computational domain of the study.

The fluid motion is governed by the incompressible Navier–Stokes equations [40]:

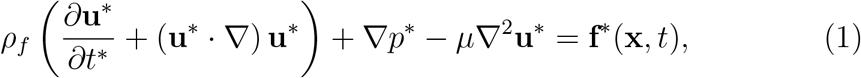

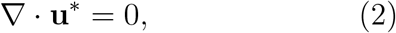

where *ρ*_*f*_ is the fluid density, **u**^*^ is the fluid velocity, *p*^*^ is the pressure, *µ* is the dynamic viscosity, and **f**^*^(**x**, *t*) represents the external force exerted by the retina on the fluid. Variables with asterisks denote dimensional quantities. The nondimensional formulation will be introduced later.

The detached retina is modeled as a one-dimensional, inextensible, inelastic filament with mass density *ρ*_*s*_ and bending stiffness *κ*_*b*_, interacting with the surrounding incompressible fluid [52]. Its motion is governed by Newton’s second law, indicating that the acceleration of the retina is directly proportional to the net force acting on it, resulting in the equation:

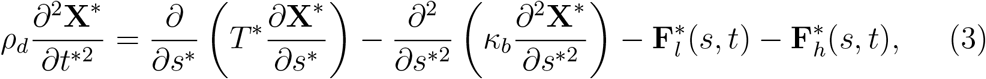

where *ρ*_*d*_ = (*ρ*_*s*_ − *ρ*_*f*_)*h*^*^ [52] is the effective areal density of the retina, *h*^*^ is the retinal thickness, *T* ^*^ is the internal tension, 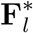 is the force exerted by the fluid on the retina, and 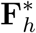 is the adhesion force restraining the retina due to bonding with the RPE.

To capture the inextensibility of the retina, we enforce the constraint:

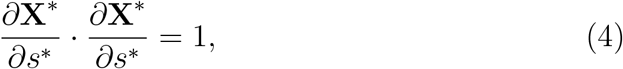

Differentiating Equation (3) with respect to *s*^*^ and taking the dot product with 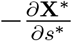 yields the equation for the internal tension:

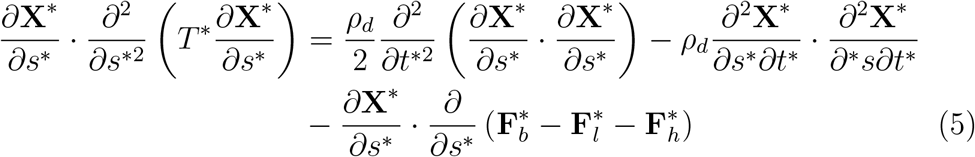

where 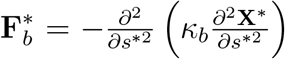 is the bending force resulting from the change in the curvature of the retina. The first term on the right hand side of Equation (5) is theoretically zero due to the inextensibility constraint but is retained to account for numerical approximation errors.

When fluid flows around a structure, both the fluid and the structure exert forces on each other. This force is reactive in nature, and its magnitude is proportional to the difference between the fluid and structure velocities, such that if the structure moves with the same velocity as the fluid, it experiences no force from the fluid. Here, the force exerted by the fluid on the retina, 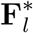, is given by Goldstein’s feedback law [31]:

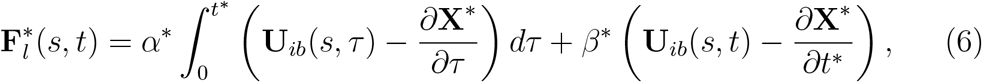

where **U**_*ib*_(*s, t*) is the interpolated fluid velocity at the fluid-structure interface:

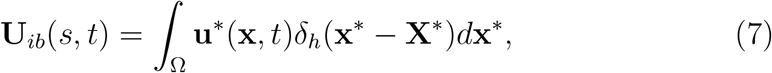

with *δ*(**x**^*^ − **X**^*^) being the two-dimensional Dirac delta function connecting the fluid’s motion to the retina. Following [54], it is given by:

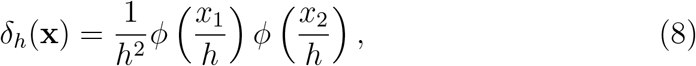

where

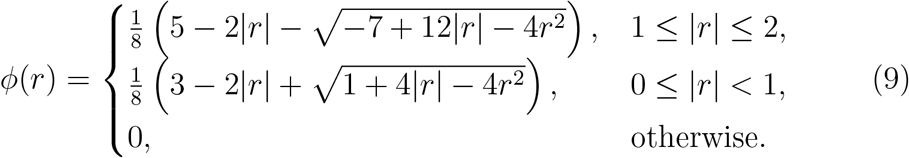

The external force density **f**^*^(**x**, *t*) that the retina exerts on the fluid is modeled as a reaction force, defined by:

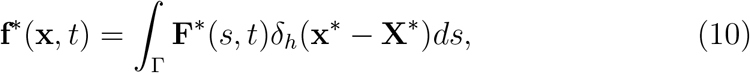

where Γ is the retina-fluid interface. This force is only experienced by fluid elements in close proximity to the retina.

### 2.2. The Adhesion Protein Model

The NL is bound to the RPE by adhesion proteins known as the interphotoreceptor matrix (IPM) [3, 37, 2, 47, 33]. To model the distribution of these proteins and the adhesion force existing between the NL and the RPE, the following assumptions are made:

1. The adhesion proteins known as IPM are uniformly distributed with density *ρ*_0_ and behave like elastic springs, offering a resistive force of *C*_*b*_ Newtons per unit length.
2. The uniform distribution *ρ*_0_ of the adhesion proteins prior to detachment becomes spatially dependent, taking values 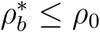 when detachment occurs.
3. The retina is more likely to peel into the vitreous chamber (i.e., move upwards in the computational domain) rather than move downwards, as the eyewall acts as a barrier. To model this directional behavior, the resistive force density offered by the adhesion proteins is defined as:

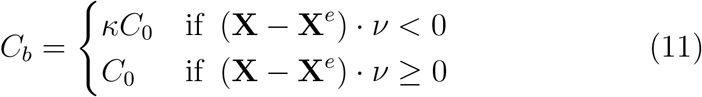

where *κ >* 1 represents magnification factor opposing downwards motion of the retina, *C*_0_ is some basal adhesion bond strength, **X**^*e*^ representing the position of the retina prior to detachment and *ν*, a unit vector perpendicular to the eyewall.
4. In regions where the retina is fully detached 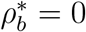, whereas in regions where the retina remains firmly attached to the RPE 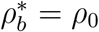. A sharp transition in adhesion density occurs near the clamped edge as shown in Figure 2.
5. In general, proteins exhibit binding and unbinding affinities[70]. We assume that the adhesion proteins (IPM) behave similarly and denote their binding and unbinding affinities by *σ*_*b*_ and *σ*_*u*_, respectively—that govern their likelihood to bind when brought into close proximity with the RPE or unbind when stretched beyond a certain threshold [67, 72].
6. Once unbound due to excessive stretch, adhesion proteins can rebind if they remain near the RPE for a sufficient time, allowing for reattachment and potential healing.
7. Binding and unbinding rates depend on the vertical separation between the NL and RPE.

Based on the above-stated assumptions, the adhesion force 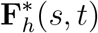, representing the restoring force between the NL and RPE, depends on the density of adhesion proteins, the strength of adhesion proteins, and the displacement of the retina from its resting configuration. The force is therefore modeled as:

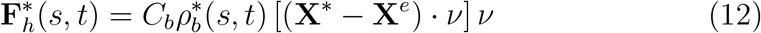

with the distribution of adhesion proteins 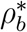 satisfying the equation:

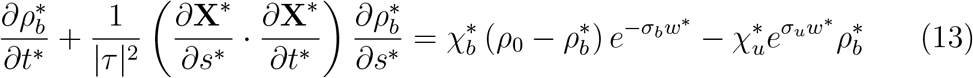

where χ_*b*_ and χ_*u*_ represent the binding and unbinding rate constants, respectively, and *w*^*^ = (**X**^*^ − **X**^*e*^) · *ν* denotes the vertical displacement of the retina. A detailed derivation of this model is presented in Appendix A.

**Figure 2:**
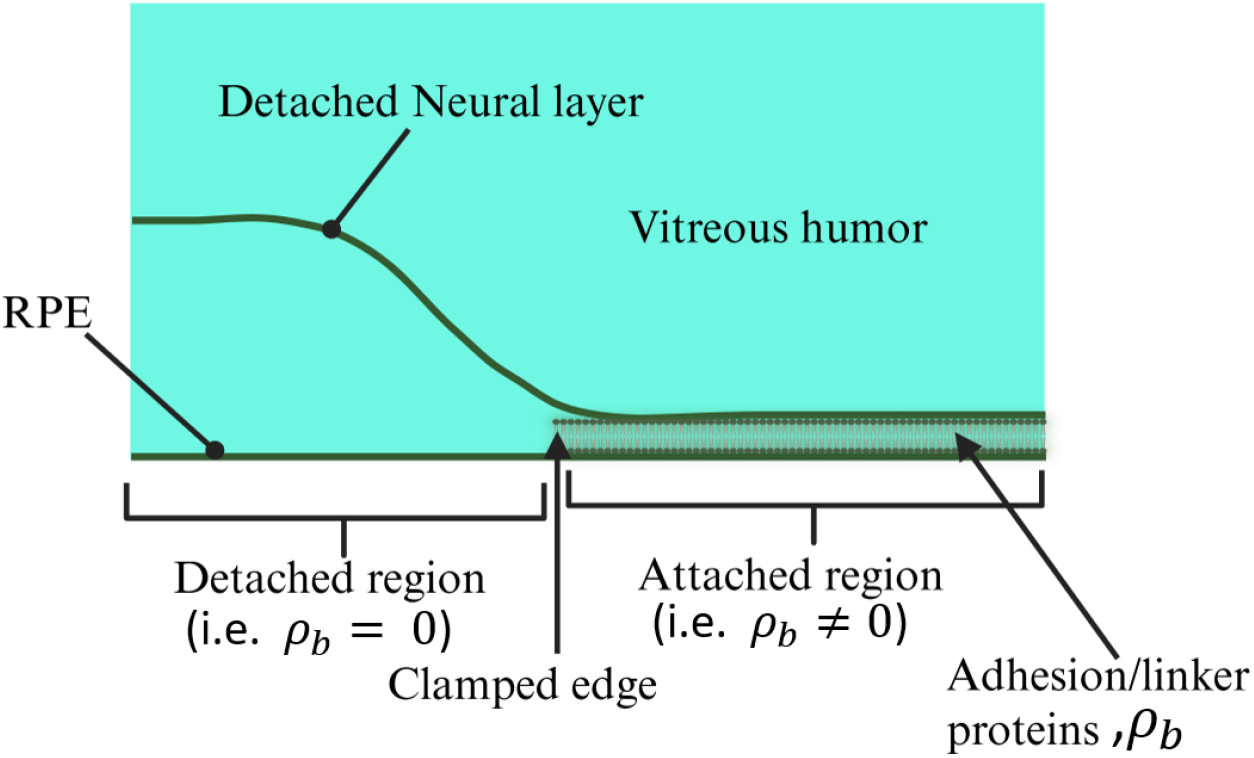
Geometry of Detachment: The figure illustrates both the detached and attached regions of the retina. It also depicts how adhesion proteins bind the neural layer (NL) to the retinal pigment epithelium (RPE).

### 2.3. Initial and Boundary Conditions

In this section, we discuss the initial and boundary conditions that must be satisfied by the RP-FSI model. Specifically, we outline the initial and boundary conditions for the fluid velocity **u**^*^, the retinal position **X**^*^, the tension force *T* ^*^, and the adhesion proteins 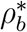.

#### 2.3.1. Initial and Boundary Conditions for the Fluid

In this work, the computational domain represents a small section of the vitreous humor, with the curvature of the eyeball neglected, as shown in Figure 1B. Since we are considering a two-dimensional flow, the fluid velocity can be expressed in component form as **u**^*^ = (*u, v*). We assume that the vitreous fluid is initially stationary and thus impose the initial condition:

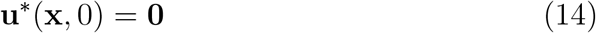

When the eyeball rotates, the vitreous fluid is set into motion due to the movement of the eyeball. Given that the fluid is viscous, we assume that fluid molecules in contact with the eyewall move along with it. Therefore, we impose a Dirichlet boundary condition

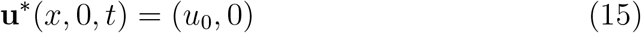

at the base of the domain.

Additionally, since the computational domain is a small section of the vitreous humor and assuming the flow is unidirectional and uniform with no turbulence, we impose a no-flux condition at the top of the domain:

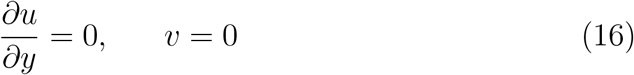

We also assume that since the computational domain is small, the rate at which fluid enters the domain on one side (either left or right) is the same as the rate at which the fluid flow out on the opposite side. To capture this, we imposed a periodic boundary conditions at the left and right boundaries:

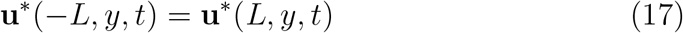

where *L* represents half the size of the domain base.

#### 2.3.2. Initial and Boundary Conditions for the Retina and the Tension

As shown in Figure 1B, and taking into account the continuous nature of the retina, the initial geometry can be reasonably approximated by a Gaussian curve defined as:

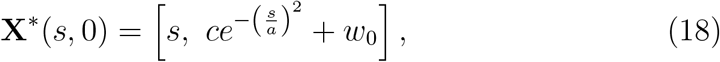

where *w*_0_ represents the resting (or natural) length of the adhesion proteins, indicating the initial vertical separation between the NL and the RPE and also containing some small amount of subretinal fluid [33]. In a healthy retina, this fluid is continuously pumped by the RPE toward the choroid, helping to maintain adhesion between the NL and the RPE [24, 16]. In defining the parameters *c* and *a* in Equation (18), we assume that adhesion bonds break when stretched beyond twice their natural length. Furthermore, we assume that the adhesion proteins at the clamped edge are already under tension, such that the initial separation between the NL and the RPE at that location is 2*w*_0_. Any additional stretching would thus result in bond breakage and detachment. Given an initial detached length *l*_0_ elevated at an angle *φ*, the arc length parameter at the clamped edge is *s*_*c*_ = *l*_0_ cos(*φ*). Equating the vertical displacement at *s* = *s*_*c*_ in Equation (18) to 2*w*_0_ and solving for *a* yields:

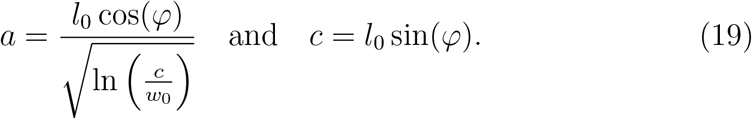

The approximation of the initial retinal geometry, highlighting the elevation angle *φ* and the role of the constant *c*, is illustrated in Figure 3.

**Figure 3:**
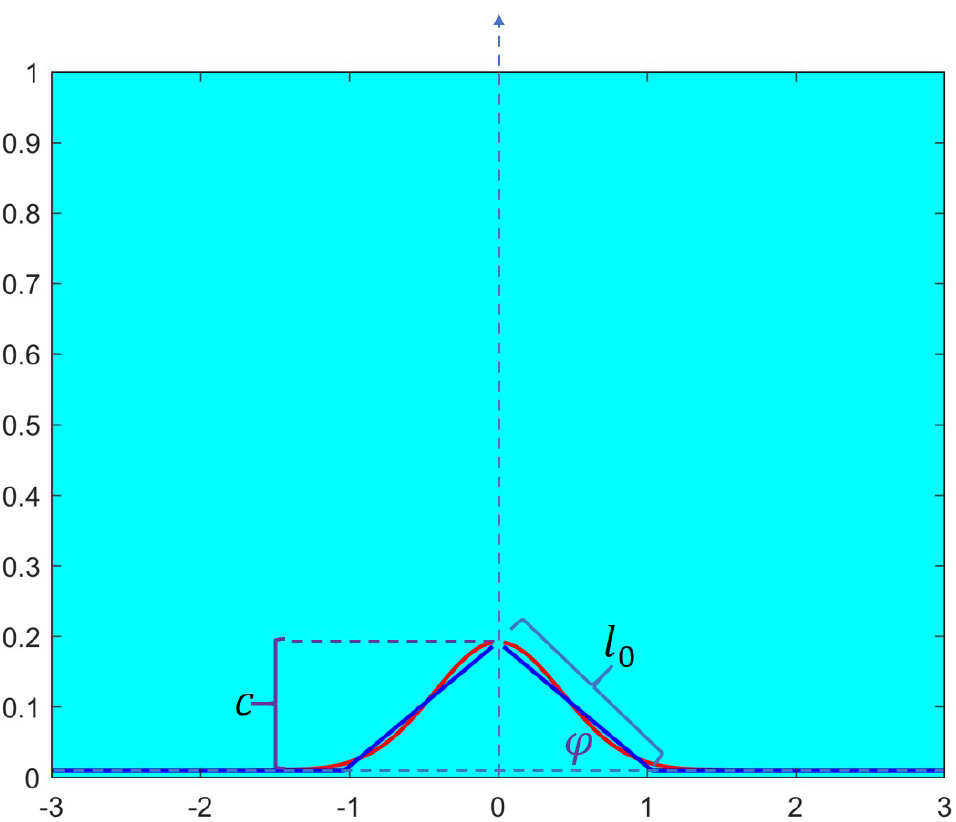
Initial Geometry of the Retina: The retina’s initial shape is approximated using a Gaussian curve, capturing the elevation and geometry of the detached segment.

At the detached end of the retina, we use the same boundary condition as Natali et. al. [52] used by assuming that both the curvature and the shear stress are zero, leading to the boundary condition:

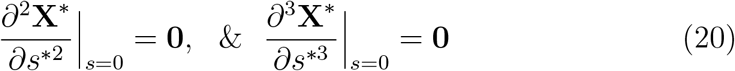

At the far end, we assume that the retina remains attached to the eyewall such that the tangent vector is horizontal. We also assume that the retina is clamped to the eyewall at this end, so that it moves with the eyewall during rotation. This gives the boundary condition:

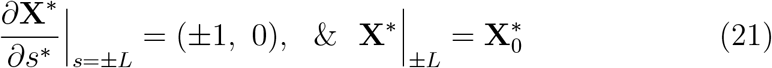

As shown in Figure 1A, in the full 3D context, both halves of the detached retina are connected. Hence, when one side is pulled, the other is dragged along. To mimic the connectivity between the two halves, we impose the boundary condition:

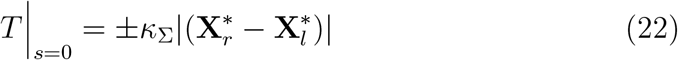

where *κ*_Σ_ is a spring constant that depends on the stiffness of the retina, and 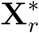 and 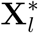 represent the right and left halves of the detached retina, as shown in Figure 1B. At the extreme ends, since the retina remains bound to the eyewall, we assume its acceleration matches that of the eyewall. Applying this condition to Equation (3) results in:

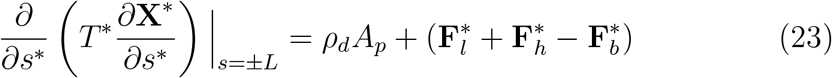

where *A*_*p*_ denotes the acceleration of the eyewall.

#### 2.3.3. Initial and Boundary Conditions for the Adhesion Proteins

As stated in the model assumptions, the distribution of adhesion proteins is zero in regions where the retina is detached. In contrast, where the retina remains fully attached, the distribution equals the pre-detachment level of available adhesion proteins. A sharp transition in the protein density occurs at the clamped edge.

The distribution of adhesion proteins governed by Equation (13) is therefore subject to the initial condition:

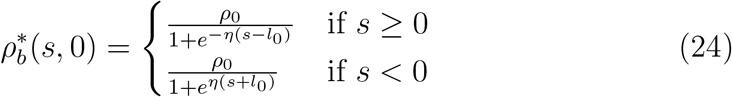

where *η* is a positive constant controlling the steepness of the transition near the clamped edge.

The initial distribution of adhesion proteins is illustrated in Figure 4, showing a zero density in the detached region and a rapid increase at the boundary where the retina becomes clamped. The system is also subject to the boundary condition: 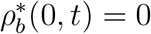

**Figure 4:**
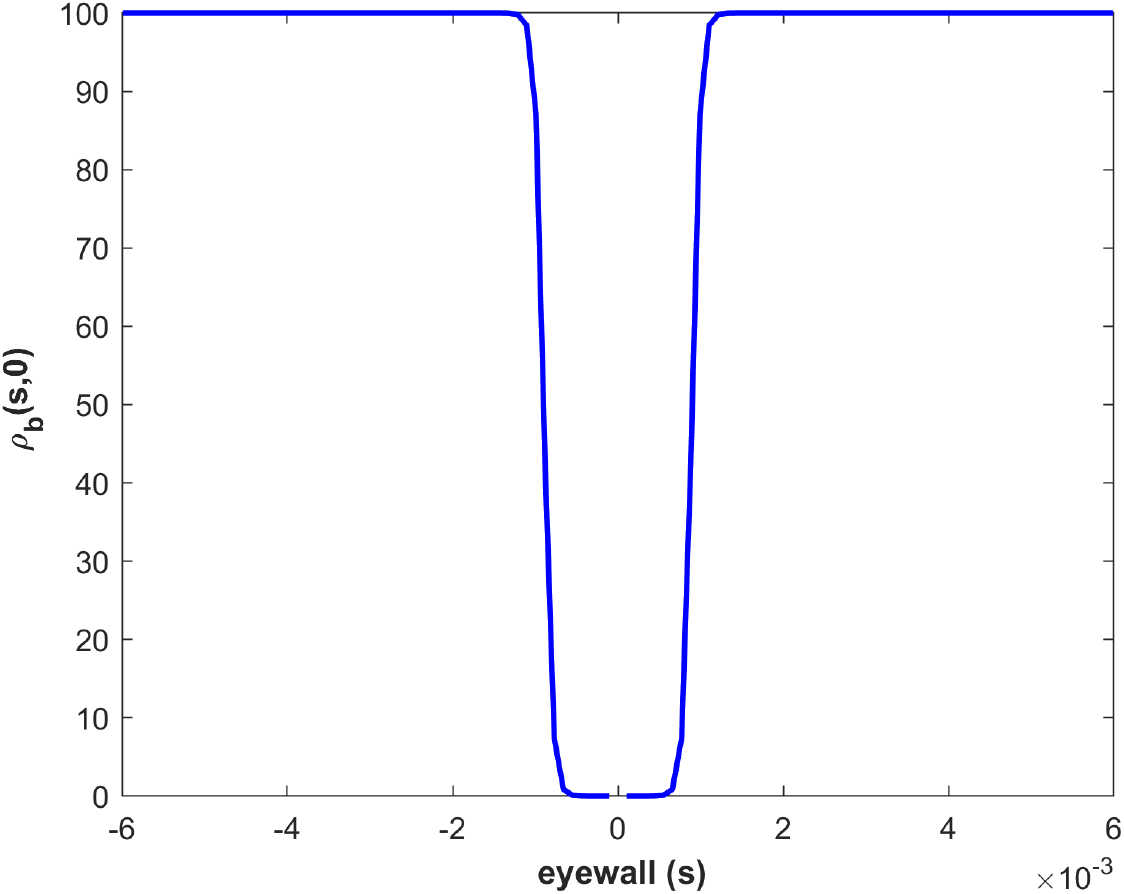
Initial distribution of adhesion proteins: The figure shows the initial profile of adhesion protein density. The density is zero in the detached region and rises sharply near the clamped edge where the retina becomes attached.

#### 2.3.4. Saccadic Eye Rotation

A saccade is a rapid, jerky movement of the eye that shifts focus from one point to another. Studies have shown that small saccadic eye rotations can be approximated using sigmoid, Gaussian, or Gumbel distribution functions [20, 56]. In this study, we modeled the time evolution of the angular displacement in degrees as the eye rotates in the horizontal direction as

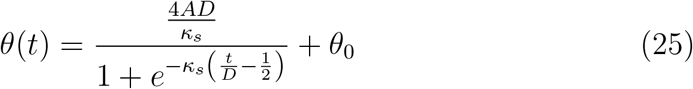

where *A* is the peak angular velocity, *D* the duration of the saccade, *θ*_0_ the initial angular displacement from the forward-looking direction, and *κ*_*s*_ a positive constant controlling the steepness of the sigmoidal curve. The angular velocity and the acceleration corresponding to the saccade are obtained by differentiating Equation (25) which are respectively given by

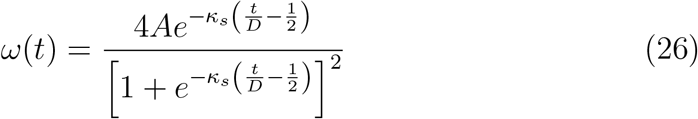

and

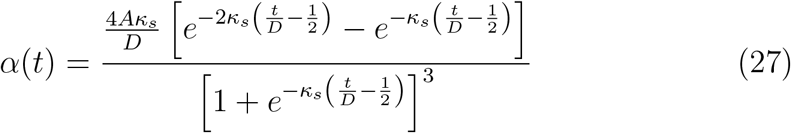

To simulate the RP-FSI model for various saccadic eye rotations, one needs to specify a sequence of peak angular velocities *A* and corresponding durations *D*. By convention, we assume *A >* 0 corresponds to rotation to the right, *A <* 0 corresponds to rotation to the left, and *A* = 0 indicates no eyeball movement.

Taking *R* as the average radius of the eye, the corresponding linear displacement, velocity, and acceleration can be computed as

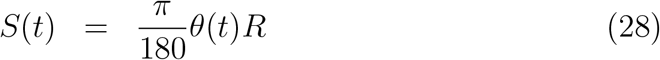

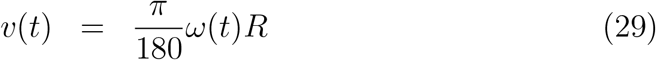

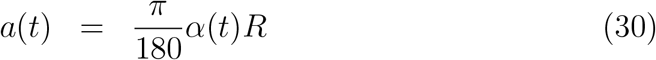

Figure 5A shows the angular displacement of the eyeball as it rotates through an angle of 8^°^ to the right and returns to the forward-looking direction. Figure 5B and Figure 5C show the corresponding linear velocity and acceleration of the eyeball.

**Figure 5:**
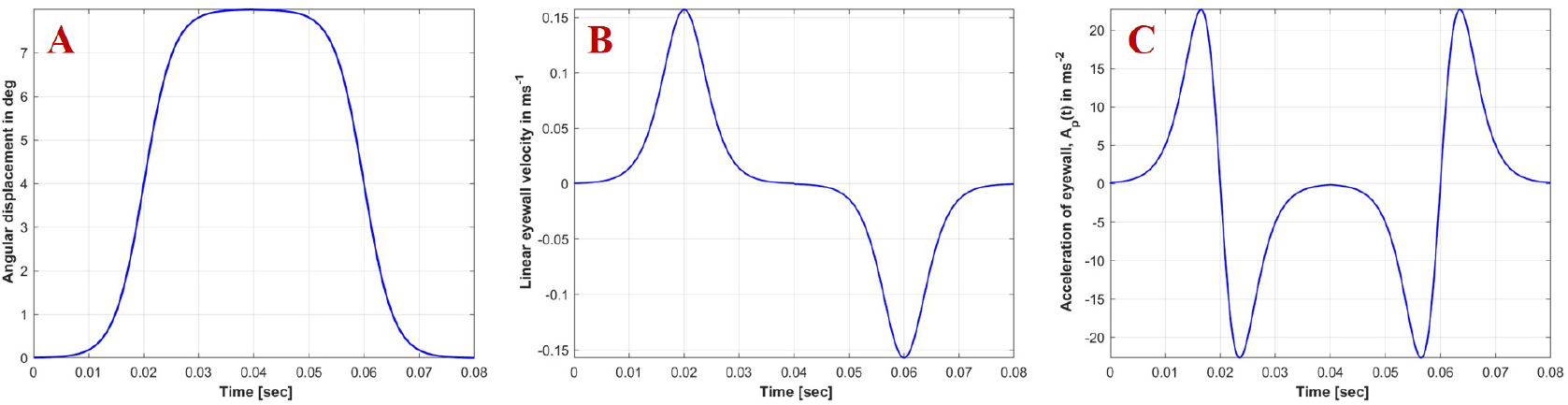
Saccadic profile: (A) Angular displacement of the eyeball during a saccadic motion, showing movement from the forward-looking direction to the right and back. (B) Corresponding linear velocity of the eyeball. (C) Corresponding linear acceleration of the eyeball.

These quantities can be nondimensionalized by scaling *S*(*t*) by a characteristic length *L*_0_ (to be discussed in the next section), *v*(*t*) by *u*_max_ = max_*t*_[*v*(*t*)], and *a*(*t*) by *a*_max_ = max_*t*_[*a*(*t*)]. The linear velocity represent the Dirichlet boundary condition specified for the fluid flow in Equation (15) whiles the linear acceleration feed into the boundary condition for the Tension as defined by Equation (23)

### 2.4. Nondimensionalization

We nondimensionalize the model presented above to reduce the number of parameters and to transform the equations into a more tractable form for analysis and numerical simulation. Dimensional quantities are denoted with asterisks, while their corresponding dimensionless forms are written without asterisks. To proceed, we introduce two characteristic parameters: *u*_*max*_, a characteristic velocity scale, and *L*_0_, a characteristic length scale. Using these, the various dimensional quantities are scaled as follows:

- The fluid pressure *p*^*^ is scaled by 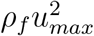
- The force density **f**^*^ exerted by the retina on the fluid is scaled by 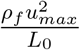
- The tension *T* ^*^ in the retina is scaled by 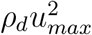
- Time *t* is scaled by 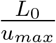
- The bending stiffness *κ*_*b*_ of the retina is scaled by 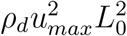
- The force 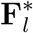 exerted by the fluid on the retina and the adhesion force 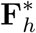 binding the NL to the RPE are scaled by 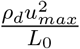
- The Goldstein feedback coefficients *α*^*^ and *β*^*^ involved in 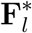 are scaled by 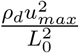 and 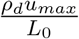, respectively
- The adhesion bond strength *C*_*b*_ is scaled by 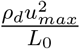
- The densities of adhesion proteins 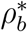 and *ρ*_0_ are scaled by 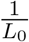
- The binding and unbinding rates of adhesion proteins, 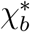 and 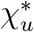, are both scaled by 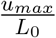

It is important to note that a two-dimensional (2D) flow can be interpreted as a special case of a three-dimensional (3D) flow in which the dynamics along the third dimension remain constant. Consequently, we retain the units of fluid and retinal densities as kg*/*m^3^. In 2D, the fluid-structure interface appears as a line; therefore, the areal mass density *ρ*_*d*_ = (*ρ*_*s*_ − *ρ*_*f*_)*h*^*^, as defined in Equation (3), is assigned the unit kg*/*m, where the second dimension is assumed to be uniform. The dimensionless form of the Navier-Stokes equations describing fluid motion becomes:

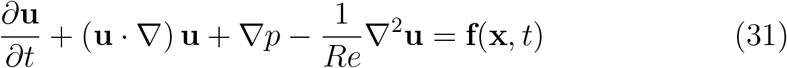

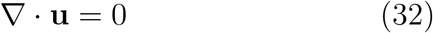

where 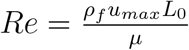 is the Reynolds number. The dimensionless form of the retina motion equation, previously presented in Equation (3), is given by:

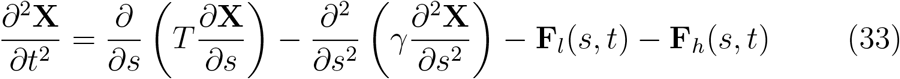

where 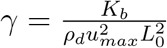 is the dimensionless bending stiffness. The dimensionless forms of the tension and inextensibility conditions in Equation (5) and Equation (4), respectively, become:

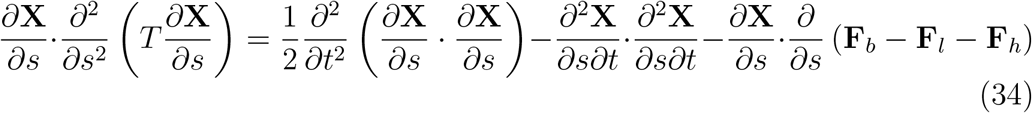

and

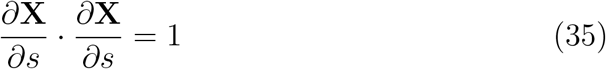

with the dimensionless forms of the interacting forces given by:

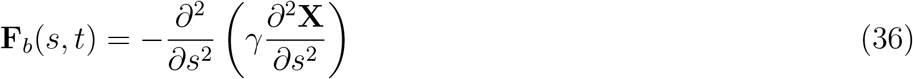

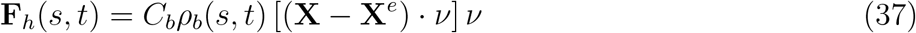

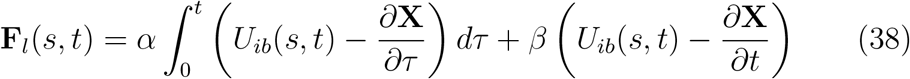

with 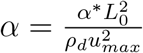 and 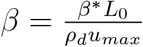 and

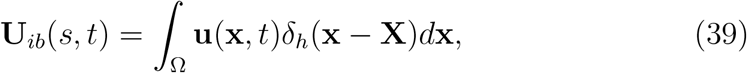

Recognizing that if **x**^*^ ∈ ℝ^*n*^, then 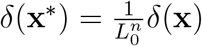 the dimensionless form of the force the retina exerts on the fluid becomes:

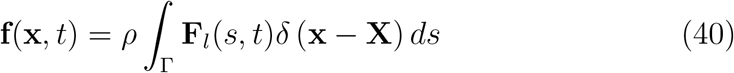

where 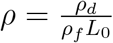.

The equation governing the density of adhesion proteins becomes:

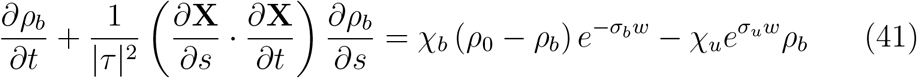

where 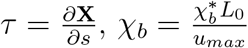 and 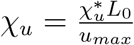.

## 3. Numerical Solution

The nondimensionalized model was solved numerically and then scaled back to dimensional form to facilitate interpretation and comparison. To simulate the RP-FSI model, the Navier–Stokes equations governing the fluid flow are solved using the non-iterative *P*_2_ projection method [8, 9]. The equation describing the adhesion protein density is solved using an implicit upwind scheme, while the model for the motion of the detached retina is solved using a finite difference approximation. In this scheme, the first term on the right-hand side of Equation (33) is treated implicitly, and the remaining terms are treated explicitly. A detailed discussion of the numerical discretizations and implementation are presented in the supplementary material. The general steps for stepping forward in time when solving the model are as follows:

1. Initialize the data by defining the initial fluid velocity **u**(**x**, 0) as given in Equation (14), the initial position of the retina **X**(*s*, 0) as defined in Equation (18), and the initial density distribution of the adhesion proteins *ρ*_*b*_(*s*, 0) as specified in Equation (24).
2. Interpolate the fluid velocity 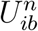 at the boundary of the retina using Equation (39) and compute the bending force 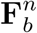(given by Equa-tion (36)), the adhesion force 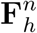 (given by Equation (37)), and the force exerted by the fluid on the retina, 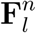 (given by Equation (38)).
3. Compute the tension 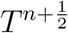 at the next half time step as defined by Equation (34), subject to the inextensibility condition in Equation (35).
4. Solve (33) to update the position of the retina, **X**^*n*+1^.
5. Compute the density 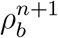 of the adhesion proteins using (41).
6. Compute the force **f**^*n*^ that the retina exerts on the fluid.
7. Solve the Navier–Stokes equations to update the velocity field **u**^*n*+1^.
8. Update the initial data for the next time step.

### 3.1. Parameter Estimation

To solve the proposed RP-FSI model, the dimensional values of the model parameters used are presented in Table 1. Some of these parameters were obtained directly from existing literature, while others were determined using related information from the literature and our model simulations.

**Table 1:** Parameter values: This table presents the baseline dimensional parameter values used in simulating the RP-FSI model. Variations around these values were explored to investigate the effects of individual parameters on detachment progression.

| Properties of the retina | Value | Reference |
| --- | --- | --- |
| Retina thickness, $h^*$ | $200 \mu m$ | [4, 49, 19, 51, 66, 60, 50] |
| Fluid viscosity, $\mu$ | $1.065 \times 10^{-3} Kg/ms$ | [61, 52] |
| Fluid density, $\rho_f$ | $1000 kg/m^3$ | [52, 17, 64] |
| Modulus of elasticity of the retina, $E$ | $1210 N/m^2$ | [28, 52] |
| Retinal density, $\rho_s$ | $1040 kg/m^3$ | [17] |
| Goldstein stiffness coefficient, $\alpha$ | $-100 Kg/ms^2$ | [52, 31] |
| Goldstein damping coefficient, $\beta$ | $-10 Kg/ms$ | [52, 31] |
| Strength of adhesion bonds, $C_b$ | $2 N/m$ | {*Estimated*} |
| Unbinding rate, $\chi_u$ | $0.6 s^{-1}$ | {*Estimated*} |
| Binding rate, $\chi_b$ | $600 s^{-1}$ | {*Estimated*} |
| Density of available adhesion proteins, $\rho_0$ | $100 m^{-1}$ | {*Estimated*} |
| Binding affinity, $\sigma_b$ | $5 \times 10^5 m^{-1}$ | {*Estimated*} |
| Unbinding affinity, $\sigma_u$ | $1.8 \times 10^5 m^{-1}$ | {*Estimated*} |
| Peak angular velocity, $A$ | $750 deg/m$ | {*Estimated*} |
| Initial detached length, $l_0$ | $0.7 mm$ | {*Estimated*} |
| Elevation of detached retina, $\varphi$ | $7^\circ$ | {*Estimated*} |
| Average radius of human eye, $R$ | $12 mm$ | [11] |
| Steepness of initial adhesion protein, $\eta$ | $2 \times 10^4 m^{-1}$ | {*Estimated*} |
| Steepness of angular displacement, $\kappa_s$ | 15 | {*Estimated*} |
| Reynold's number, | $Re = \frac{\rho_f u_0 L_0}{\mu}$ | |
| Bending rigidity of the retina | $\kappa_b = \frac{E h^*{}^3}{12} Nm^2$ | [52] |
| strength at the tip connection | $K_\Sigma = \kappa_b$ | |

The computational domain was chosen as a rectangular region measuring 12 × 2 mm, with the fluid height above the eyewall set to 2 mm. The initial detached length for each half of the retina (left and right) was set to *l*_0_ = 0.7 mm, representing 11.67% of half the domain length. In the absence of precise data for the vertical elevation of the detached retina, we assumed a small elevation and selected an angle of 7^°^. The characteristics length *L*_0_ used to scaled all length related parameters was chosen to correspond to the height of the fluid above the eyewall.

A study by Henri deGuillebon et al. [22] estimated that a force of approximately 180 mN*/*m is required to separate the NL from the RPE in a rabbit eye. They also observed that the adhesion strength varies significantly with location. Based on our formulation of the adhesion force presented in Equation (12), we explored appropriate values for *C*_*b*_, starting with the average force reported by Henri deGuillebon et al. The value of *C*_*b*_ presented in Table 1 was selected after running multiple simulations and analyzing the resulting detachment profiles.

The source term in our model for the distribution of adhesion bonds (given by Equation (13)) is similar in structure to the one used by Alert et al. [5] in their study of bleb formation dynamics. Although our model addresses a different biological process, we begun simulations with the binding and unbinding rates reported in their study. Through iterative simulations and observing the detachment profile, we narrowed down and arrived at suitable values for the binding and unbinding rates *χ*_*b*_ *& χ*_*u*_ listed in Table 1.

Due to the lack of experimental data for the density of available adhesion proteins *ρ*_0_, as well as the binding (*σ*_*b*_) and unbinding (*σ*_*u*_) affinities, we performed multiple simulations in which these parameters were varied independently. Parameter values were selected based on the resulting adhesion dynamics, ensuring alignment with physiologically plausible behavior (i.e further detachment of the retina). For the eye rotation, we adopted a peak angular velocity that results in an 8^°^ eye movement within 0.04 seconds, following a rough estimate of saccadic eye movement duration reported by Natali et al. [52]. Based on preliminary explorations, we selected the parameter *κ*_*s*_ to yield an appropriate steepness in the displacement profile. The parameter *η* in Equation (24), which controls the steepness of the adhesion protein density near the clamped edges, was chosen to produce a suitable rise at the clamped edge, as illustrated in Figure 4.

## 4. Results and Discussion

In this section, we present numerical results from the simulation of our proposed RP-FSI model developed to study the progression of rhegmatogenous retinal detachment (RRD). We consider a complete eye rotation consisting of two saccadic movements: the eyeball first rotates to the right and then returns to its forward-looking position. Each saccade is assumed to last 0.04 sec, resulting in a total simulation time of 0.08 sec.

We begin by presenting results that verify the convergence of the numerical scheme used to solve the RP-FSI model. We examine convergence of the discretization of fluid velocity and retinal position in space and time. Next, we present simulation results obtained using the base parameter values listed in Table 1. These include visualizations of the flow field interacting with the retina and the distribution of adhesion bond density, highlighting how the adhesion bonds are broken resulting in detachment progression as the fluid interacts with the detached retina during eye rotation.

Subsequently, we investigate the influence of individual model parameters on the detachment progression. In this parametric study, all other parameters are held constant while one parameter is varied at a time. By analyzing how this effective detached length evolves over the course of one complete eye rotation, we gain insight into how different physical, biological, and geometric properties—such as those of the fluid, retina, and adhesion proteins—influence the dynamics of retinal detachment.

### 4.1. Convergence Analysis

The *P*_2_ projection method used to solve the Navier–Stokes equations is formally second-order accurate, whereas the implicit upwind scheme is first-order accurate. Thus, the combination of these two schemes is expected to result in an overall convergence order between first and second. To assess the convergence of the numerical method, we introduce *û*as a refined approximation of the exact solution, computed using either a very small time step or spatial resolution. Since a closed-form analytical solution is unavailable, this refined numerical solution is used as the true solution.

To assess convergence in space (respectively, time), we fixed the time step and varied the spatial resolution. We then computed the Frobenius norm of the error between the solution at each spatial step and the refined solution *û*. Let *h* denote the spatial step size, and define *u*(*h*) as the approximate solution computed using step size *h*. Assuming the numerical scheme has order *p* and *h* is sufficiently small, the numerical errors at step sizes *h* and *h/*2 are given by:

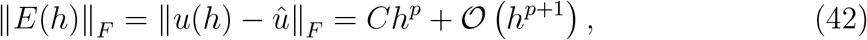

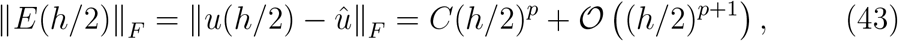

for some constant *C* independent of *h*. Dividing Equation (42) by Equation (43) and solving for *p*, we obtain the order of convergence *p* as:

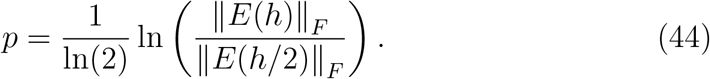

Since the fluid is defined in 2D, assembling all the *u* and *v* components of the fluid velocity **u** over the computational domain results in velocity component-wise matrices of dimensions (*N*_*y*_ + 2) × (*N*_*x*_ + 1) and (*N*_*y*_ + 1) × (*N*_*x*_ + 2), respectively, where *N*_*x*_ and *N*_*y*_ denote the number of computational cells in the *x* and *y* directions. In the staggered grid representation, the *u* and *v* components of the fluid velocity are located at different spatial positions (i.e., along the vertical and horizontal edges of the computational cells, respectively). Thus, the fluid velocity can be regarded as the union of these two matrices. To compute the error in the fluid velocity, we use the Frobenius norm, defined as:

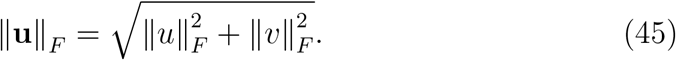

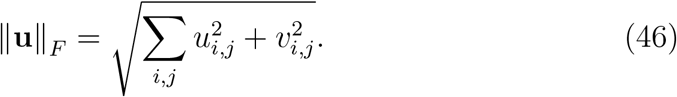

#### Convergence in space

To obtain convergence in space, we fixed the time step at *dt* = 1 × 10^−5^ and computed the reference solution using a spatial step of *h* = 6.25 ×10^−5^. The same spatial resolution was used for both the fluid and the retinal equations. Table 2 illustrates the error and the order of convergence for both the fluid velocity and the position of the retina as the spatial step is varied. We observed that the fluid velocity **u** converges with an approximate order of 1.3949, while the retina position **X** converges with an order of 1.9764. Figure 6A and B respectively show the plots of the error in the fluid velocity and the retinal position as the spatial step is varied and the corresponding approximate solutions compared to the refined solution.

**Table 2:** Table of Spatial Convergence Order: This table presents the numerical errors and the corresponding order of convergence with respect to spatial resolution.

| Spatial step, $h$ | Error in $u$ | Order in $u$ | Error in $X$ | Order in $X$ |
| --- | --- | --- | --- | --- |
| 0.001 | 0.24666 | - | $6.5274 \times 10^{-3}$ | - |
| 0.0005 | 0.19909 | 0.30912 | $2.98 \times 10^{-3}$ | 1.2637 |
| 0.00025 | 0.087767 | 1.1817 | $1.206 \times 10^{-3}$ | 1.3614 |
| 0.000125 | 0.033376 | 1.3949 | $0.53193 \times 10^{-3}$ | 1.9764 |

**Figure 6:**
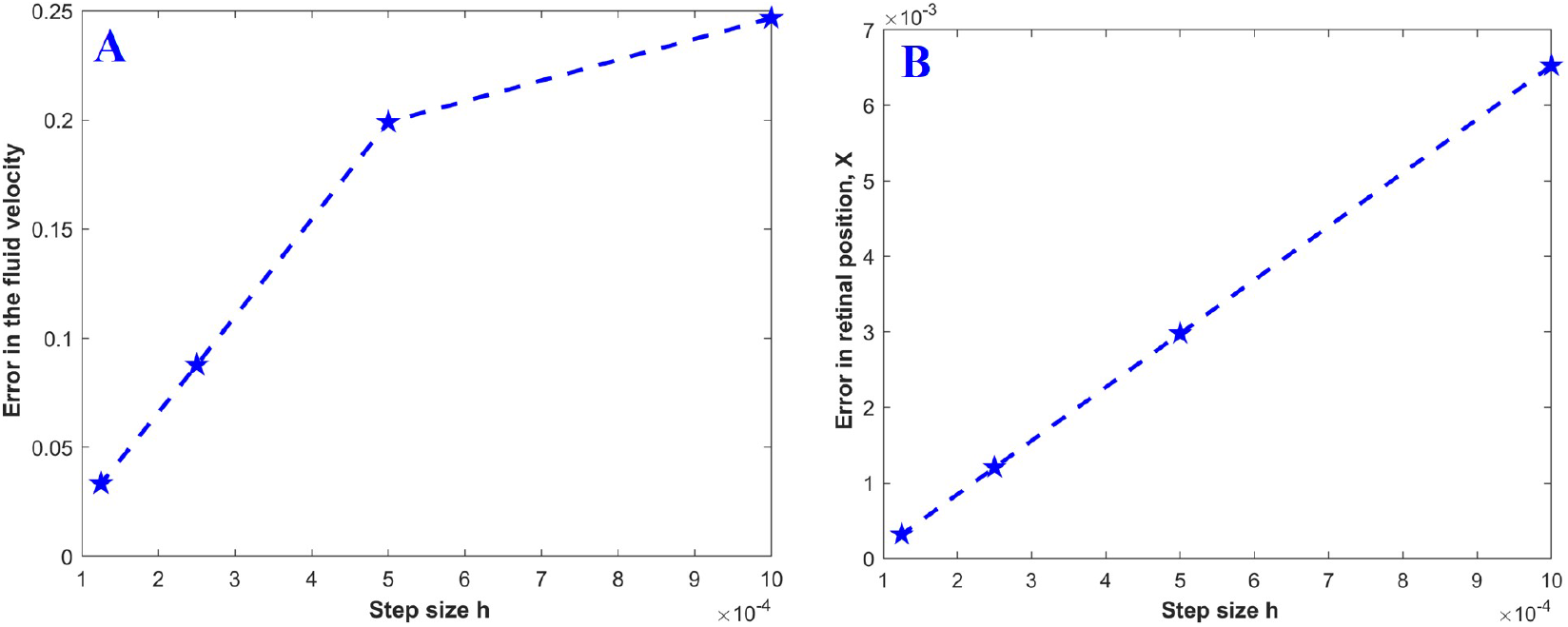
Convergence in space: (A) shows the convergence in space for the fluid velocity, while (B) shows the convergence in space for the position of the retina.

#### Convergence in time

Here, we fixed the spatial step at *h* = 5 × 10^−5^ for both the fluid velocity and the retinal position. The refined solution was computed using a time step of 1.25 × 10^−6^. Then, by varying the time steps, we computed approximate solutions for both the fluid velocity **u** and the retinal position **X** and compared them with the refined solution.

Table 3 shows the error and the approximate order of convergence as the time steps are varied, while Figure 7A and B respectively show the plots of the error with varying time steps. We observe that the fluid velocity converges with an approximate order of 1.5671, while the retinal position converges with an approximate order of 0.61575. The time steps used to compute the approximate solutions were generally small, resulting in very low numerical errors when compared with the refined solutions, as shown in Figure 7B. This could explain why the order of convergence for the retinal position was less than one.

**Table 3:** Table of Temporal Convergence Order: This table presents the numerical errors and the corresponding order of convergence with respect to temporal resolution.

| Time step, $dt$ | Error in $u$ | Order in $u$ | Error in $X$ | Order in $X$ |
| --- | --- | --- | --- | --- |
| $4 \times 10^{-5}$ | $9.1822 \times 10^{-4}$ | - | $2.3712 \times 10^{-7}$ | - |
| $2 \times 10^{-5}$ | $4.4425 \times 10^{-4}$ | 1.0475 | $2.0327 \times 10^{-7}$ | 0.22221 |
| $1 \times 10^{-5}$ | $2.0777 \times 10^{-4}$ | 1.0964 | $1.7536 \times 10^{-7}$ | 0.21311 |
| $0.5 \times 10^{-5}$ | $0.89551 \times 10^{-4}$ | 1.2142 | $1.4501 \times 10^{-7}$ | 0.27415 |
| $0.25 \times 10^{-5}$ | $0.30223 \times 10^{-4}$ | 1.5671 | $0.94632 \times 10^{-7}$ | 0.61575 |

**Figure 7:**
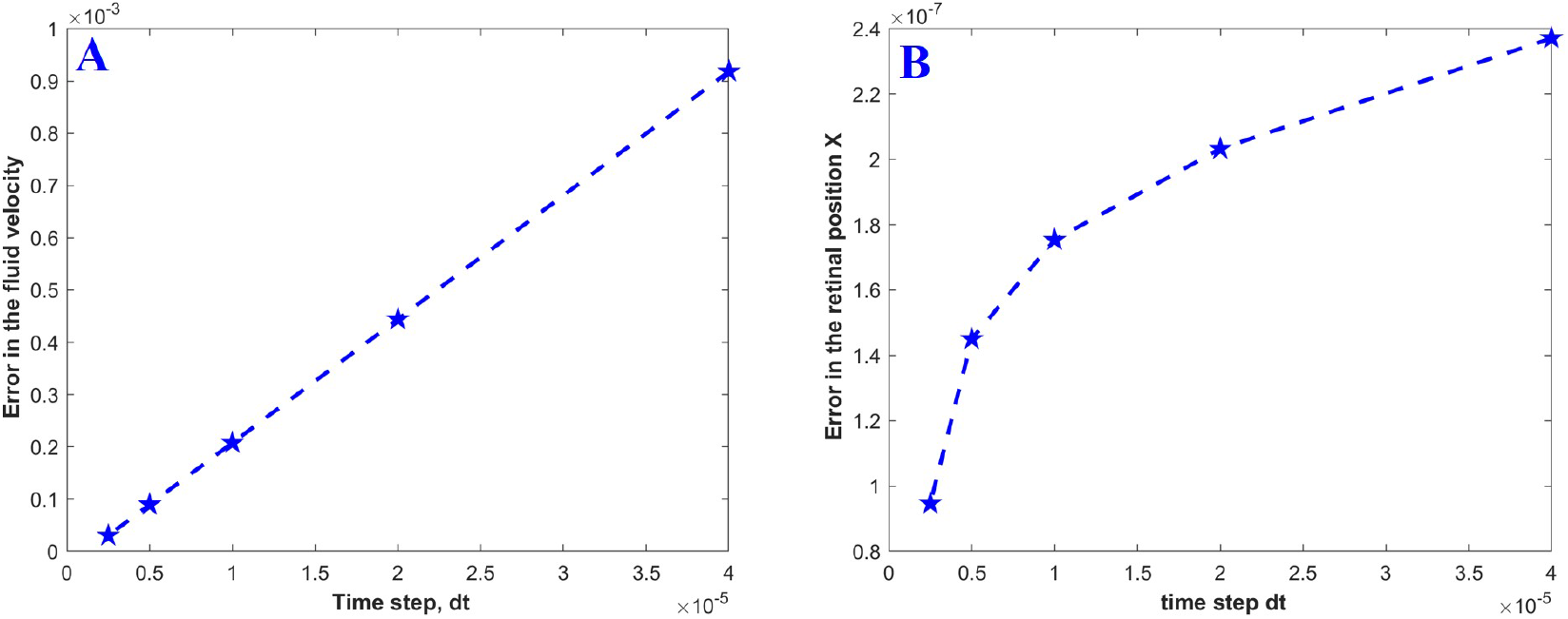
Convergence in time: (A) shows the convergence in time for the fluid velocity, while (B) shows the convergence in time for the position of the retina.

In all, these results confirm that the overall numerical scheme achieves an order of convergence between first and second order, as expected due to the combination of second-order (*P*_2_ projection) and first-order (implicit upwind) schemes used in solving the model.

### 4.2. Results from baseline parameters

Using the baseline parameter values presented in Table 1, we examine how the fluid interacts with the retina and how the adhesion bonds are broken, resulting in further detachment. We also visualize the profile of the peeling as the eyeball rotates and the fluid interacts with the retina. This is done by examining the case of a complete eye rotation, in which the eyeball first rotates to the right before returning to the forward-looking position.

Figure 8 presents snapshots of the flow field over time, where the background color represents the magnitude of the fluid velocity. The initial geometry of the retina is shown in black, while its positions at subsequent time points are shown in red. The velocity profile of the eyewall, shown in Figure 5B, indicates that the eyewall attains its peak velocity at *t* = 0.02 sec and *t* = 0.06 sec when it is moving to the right and returning to the left, respectively. This is clearly reflected in Figure 8, where the magnitude of the fluid velocity is high near the eyewall at those times.

**Figure 8:**
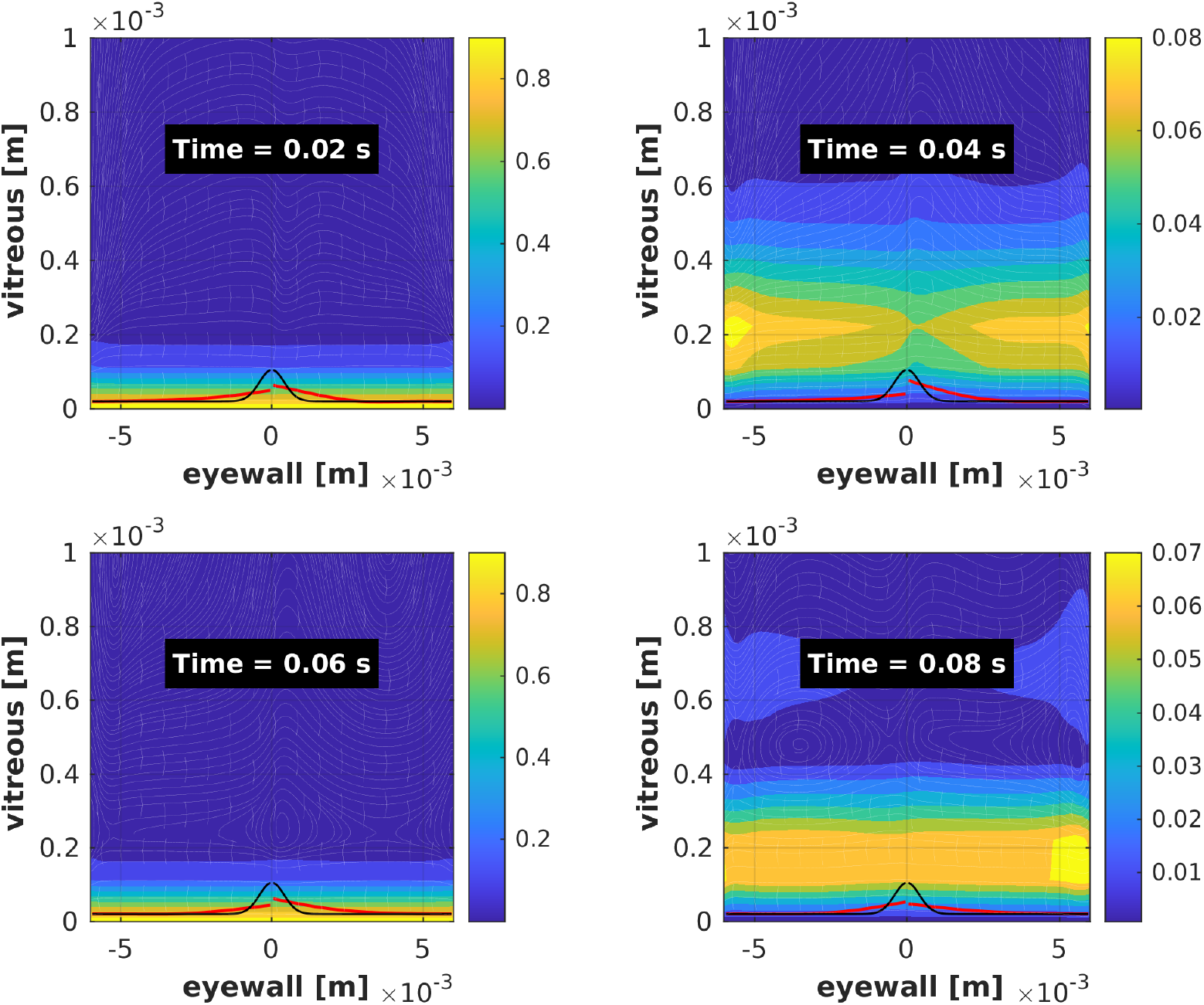
Snapshot of flow profile field: Snapshots of the fluid velocity profile and retina position at different time points using the baseline parameter values. Background color indicates the magnitude of the velocity field.

At *t* = 0.04 sec, a transition occurs that leads to a change in flow direction. Here, the fluid velocity near the eyewall is low, but high away from the eyewall, allowing the fluid molecules near the eyewall to reorient as the eyeball changes direction. At *t* = 0.08 sec, the eyewall stops moving, but the motion of the fluid does not immediately cease. This results in a low-velocity field near the eyewall and a high-velocity field away from it. The change in the position of the retina as the fluid flows demonstrates that the interaction between the fluid and the retina results in further detachment.

To visualize this detachment progression, we examine the density profile of the adhesion proteins at different time snapshots. As the detachment progresses, the density distribution of adhesion proteins also evolves. In regions where the retina is fully detached, the adhesion protein density *ρ*_*b*_ becomes zero. A sharp increase in *ρ*_*b*_ is observed at the clamped edge, which helps identify the detachment front. Specifically, the detached length can be computed as the distance from the origin to the point where *ρ*_*b*_ begins to rise. Figure 9 shows snapshots of the adhesion protein distribution over time, highlighting the evolution of detachment. We observed that detachment progressed more rapidly during rightward eye rotation, with reduced progression upon reversal, likely because energy is first used to reorient the fluid and the retina initially moving rightward. This therefore reduces the impact of the flow on the retina.

**Figure 9:**
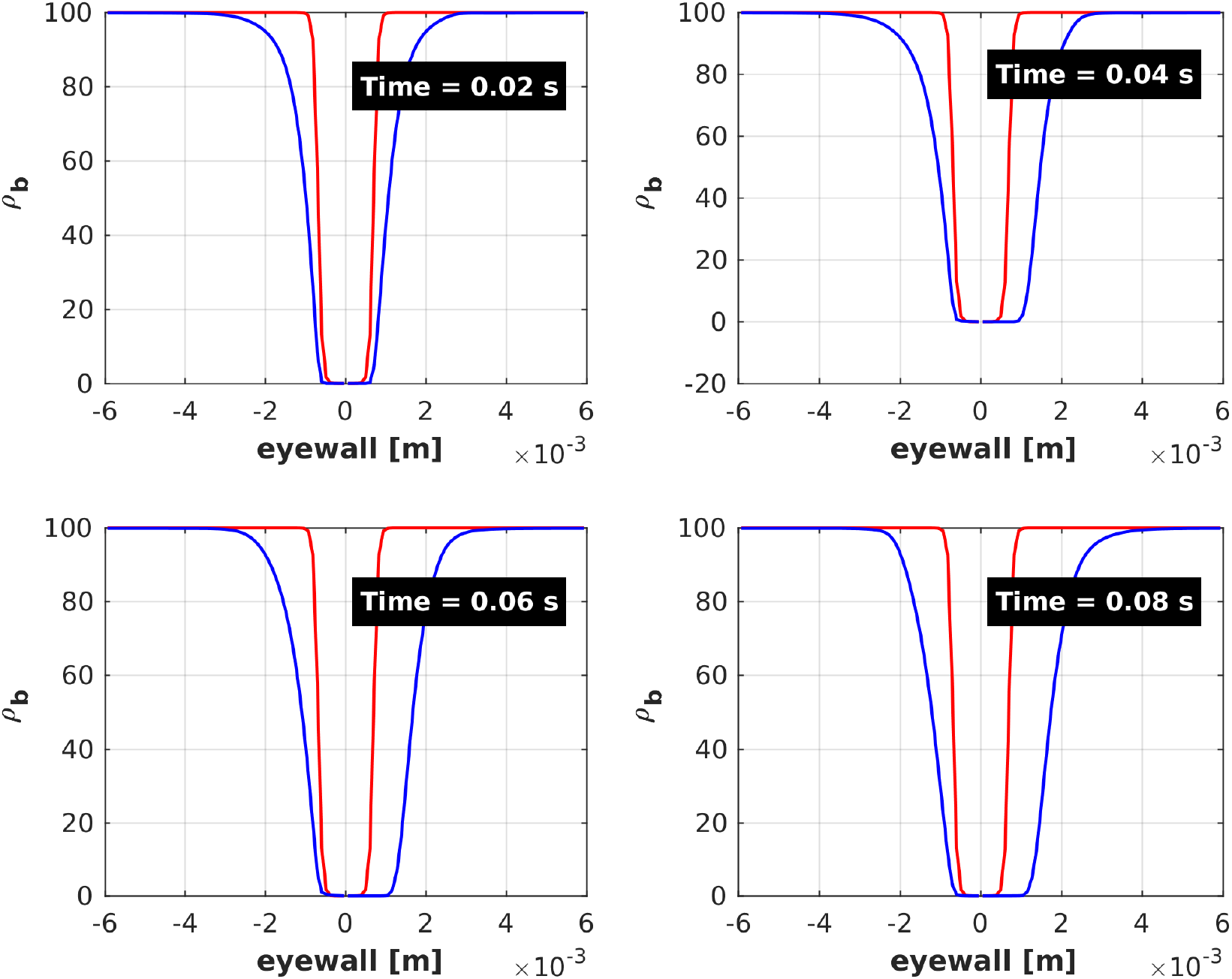
Snapshot of adhesion density profile: The figure shows snapshots of the adhesion protein density at different time points using the baseline parameter values. The horizontal distance of the blue line from the origin indicates the extent of retinal detachment.

The gradual detachment of the retina—represented by the increase in the detached length, defined as the distance from the origin to the clamped edge—is illustrated in Figure 10. We observed that as the fluid interacts with the detached retina during eye rotation, the detached length gradually increases over time. This figure provides a clear visualization of how detachment evolves continuously over the 0.08 sec simulation period as the eye rotates.

**Figure 10:**
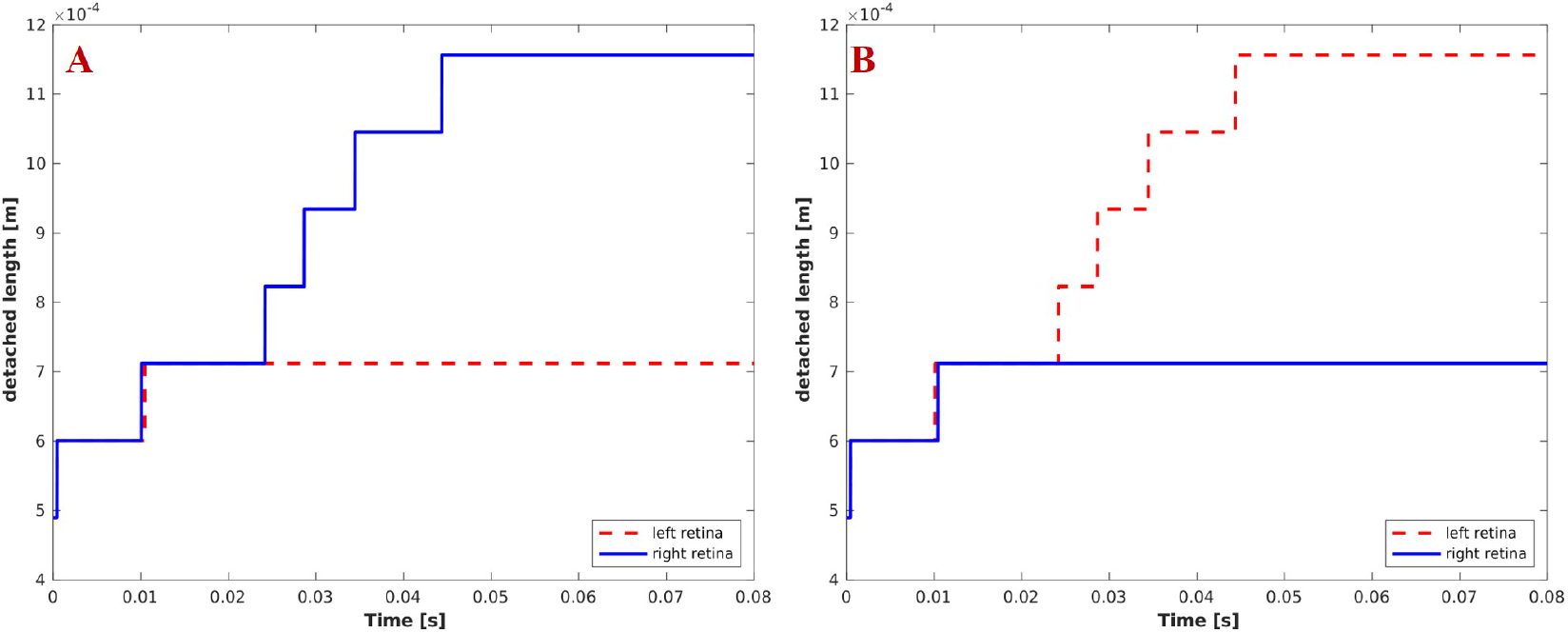
Retinal detachment profile: The figure shows the progression of retinal detachment over time. The detached length is measured from the retinal hole to the clamped edge, where *ρ*_*b*_ *>* 0, as defined in Figure 9. Panel (A) corresponds to an initial rightward rotation, and panel (B) to an initial leftward rotation.

We observed that when the eye first rotates to the right, there is more detachment in the right half of the retina compared to the left, as shown in Figure 10A. To further investigate the effect of the initial direction of eye rotation on the detachment progression, we simulated the model for the case where the eyeball first rotated to the left. The results, shown in Figure 10B, indicate that there is more detachment in the left half of the retina compared to the right. This non-symmetric behavior occurs because, when the eyeball initially rotates to the right, the fluid is propelled in the same direction as the eyewall. When the eye reverses direction, some energy is required to decelerate and then redirect the fluid flow to the left, resulting in a diminished impact on the left half of the retina.

For the remainder of the analysis, we focus on the case where the eye initially rotates to the right. All further assessments of effective detachment—defined as the distance between the final and initial detached lengths—will be based on the detachment observed in the right half of the retina, with the understanding that similar, though slightly less pronounced, detachment occurs on the left side.

### 4.3. Previously Studied Parameters

Natali et al. [52] developed a fluid–structure interaction (FSI) model to investigate the propensity for further detachment at the clamped edge of the retina. Although they did not compute the actual detached length, their analysis revealed that increasing the initial detached length, the elevation angle during an 8^°^ eye rotation, and the retinal density (*ρ*_*s*_) all contribute to an increased likelihood of further detachment at the clamped edge. In our simulation, we computed the effective detached length, defined as the distance between the final and initial detached lengths that can occur during one complete eye rotation.

Consistent with their findings, our simulations also show that increasing any of these three parameters leads to a larger effective detached length. Figure 11A and B respectively illustrate how the effective detached length evolves when either the initial detachment length (*l*_0_) or the elevation angle (*φ*) is varied independently, while keeping all other parameters fixed. The possible explanation of this results is that increasing the initial detachment or the elevation angle exposes a larger surface area of the retina to the fluid flow, thereby amplifying the fluid’s impact and leading to greater detachment.

**Figure 11:**
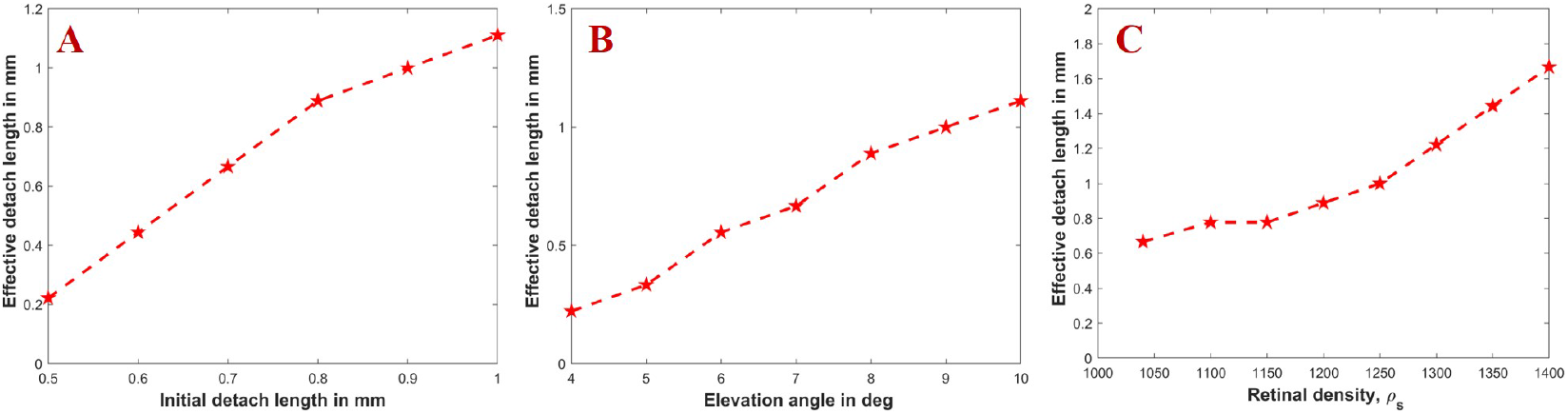
Effects of initial detached length, elevation angle, and retinal density: (A) illustrates the effect of increasing the initial detached length on the effective detachment; (B) shows how increasing the elevation angle at the clamped edge of the detached retina influences the effective detached length; and (C) demonstrates how varying the retinal density affects the effective detached length.

The inertial force acting on the retina, as shown in Equation (3), is influenced by the areal mass density, which is proportional to the difference between the retinal and fluid densities. Increasing the retinal density while keeping the fluid density constant leads to a higher areal mass density and consequently a greater inertial force on the retina. This results in increased detachment, as demonstrated in Figure 11C. These observations suggest that the closer the retinal density is to that of the fluid, the slower the detachment progression.

### 4.4. Unexplored Model Parameters

In this subsection, we examine how additional model parameters—describing fluid, retinal, and molecular properties—influence the effective detached length during one complete eye rotation. Natali et al. [52], in their work, introduced parameters such as retinal thickness *h*^*^, elasticity of the retina denoted by *E*, dynamic viscosity of the fluid denoted by *µ*, the fluid density *ρ*_*f*_, and the Goldstein feedback coefficients *α & β* defining the force the fluid exert on the retina, but did not investigate how changing these parameters could affect the tendency to detach.

#### 4.4.1. Effect of Fluid and Retinal Properties

Studies have shown that, with age, the vitreous humor gradually loses its gel-like nature and becomes more liquefied [57, 61]. To assess how this transition affects detachment, we investigate how changes in the viscosity and density of the vitreous humor influence the effective detached length during one eye rotation. As shown in Figure 12A and B, variations in fluid viscosity has relatively little effect on the progression of detachment. However, a decrease in fluid density results in a noticeable increase in the effective detached length.

**Figure 12:**
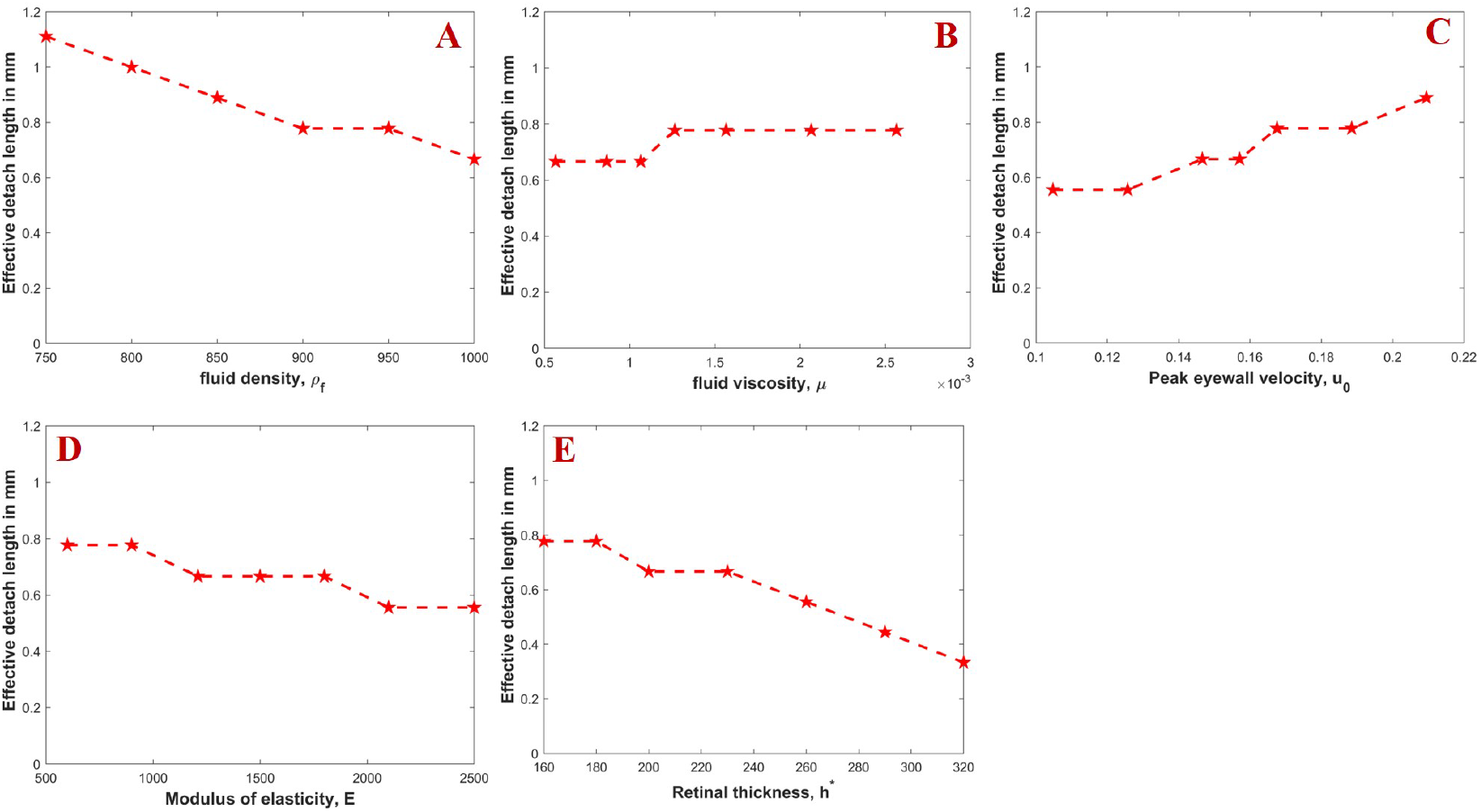
Fluid and retinal properties: (A) shows the effect of varying fluid density on the effective detached length; (B) illustrates how increasing the dynamic viscosity of the fluid influences detachment progression; (C) shows the impact of eye rotation speed on the effective detached length; (D) and (E) respectively show how retinal elasticity and thickness affect the progression of detachment

From a modeling perspective, as shown in Equation (3), the motion of the retina is influenced by the areal mass density *ρ*_*d*_ = (*ρ*_*s*_ − *ρ*_*f*_)*h*^*^. This suggests that reducing the fluid density increases the areal mass density, which in turn increases the inertial force acting on the retina, thereby resulting in a higher effective detached length. A physical explanation is that less dense fluids offer less resistance (known as drag) to the motion of immersed structures, so with the same amount of fluid velocity, the retina is able to move faster when the density is low, resulting in greater detachment.

Viscosity generally measures a fluid’s internal resistance to flow, such that increasing viscosity reduces fluid velocity. Even though the velocity of the fluid may reduce, the force the fluid exerts on the retina, as defined by the Goldstein feedback law in Equation (6), can still remain high since it depends on the difference between the fluid and retinal velocities. This may explain why we observe less change in the effective detached length as the viscosity is varied. The highly nonlinear coupling of the model makes it not obvious to clearly explain, from a modeling perspective, how precisely the viscosity affects the detachment progression.

Several studies have identified a correlation between increasing age and the risk of retinal detachment, with older individuals exhibiting a higher likelihood of developing the condition [43, 21, 27]. It has also been observed that both retinal thickness [39] and elasticity [45, 14] decrease with age. However, to the best of our knowledge, no prior studies have investigated how these changes influence the progression of detachment.

From the modeling perspective, the bending stiffness 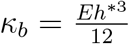, as pre-sented in Table 1, depends on the product of the modulus of elasticity *E* and the cube of the retinal thickness *h*^*^. From Equation (3), we observe that increasing the bending stiffness reduces the net force acting on the retina. In a biological context, increasing the thickness and elasticity of the retina enhances its ability to withstand the pressure exerted by the fluid, thereby slowing down the progression of detachment. Our model simulations, shown in Figure 12D and E, demonstrate that increasing either the retinal density or thickness leads to a reduction in effective detached length. This suggests that individuals with thinner and less elastic retinas (i.e., typically older individuals) are not only more susceptible to the onset of detachment but also to its faster progression.

The progression of retinal detachment is primarily driven by the motion of the eyeball, which sets the vitreous humor into motion and generates fluid forces that push against the detached retina. To further explore this mechanism, we investigate how the speed of eye rotation affects detachment.

As illustrated in Figure 12C, increasing the speed of eye rotation results in greater effective detachment. From the boundary condition for the fluid flow given by Equation (14), we imposed a Dirichlet boundary condition **u** = (*u*_0_, 0) at the base of the computational domain, where *u*_0_ represents the velocity of the eyewall. Since the fluid is viscous, we assume that fluid molecules in contact with the eyewall move at the same speed as the eyewall. Therefore, increasing the velocity of eye rotation leads to greater movement of the fluid molecules, which in turn increases the force exerted on the retina, thereby enhancing the detachment. This results suggests that patients with retinal detachment may benefit from minimizing rapid eye movements to slow the disease’s progression.

#### 4.4.2. Molecular Properties

The NL is attached to the RPE by the interphotoreceptor matrix (IPM) [3, 37, 2, 47, 33], which we refer to as adhesion proteins. Proteins in general (including IPM) exhibit both binding and unbinding affinities [67, 72], which determine their likelihood of attaching to or detaching from a binding site when brought into close proximity or subjected to stress. Additionally, different proteins form bonds of varying strength. In this section, we examine how molecular properties—including adhesion bond density, bond strength, binding/unbinding affinities, and binding/unbinding rates—affect the progression of retinal detachment.

When the number of adhesion proteins or the strength of the bond these proteins form increases, the NL become firmly bound to the RPE thereby making it difficult to separate. From Figure 13A and B, we observe that increasing either the density or the strength of adhesion bonds slows down the rate of progression. This suggests that the stability of the retina’s attachment to the RPE depends, in part, on how firmly the two layers are bonded.

**Figure 13:**
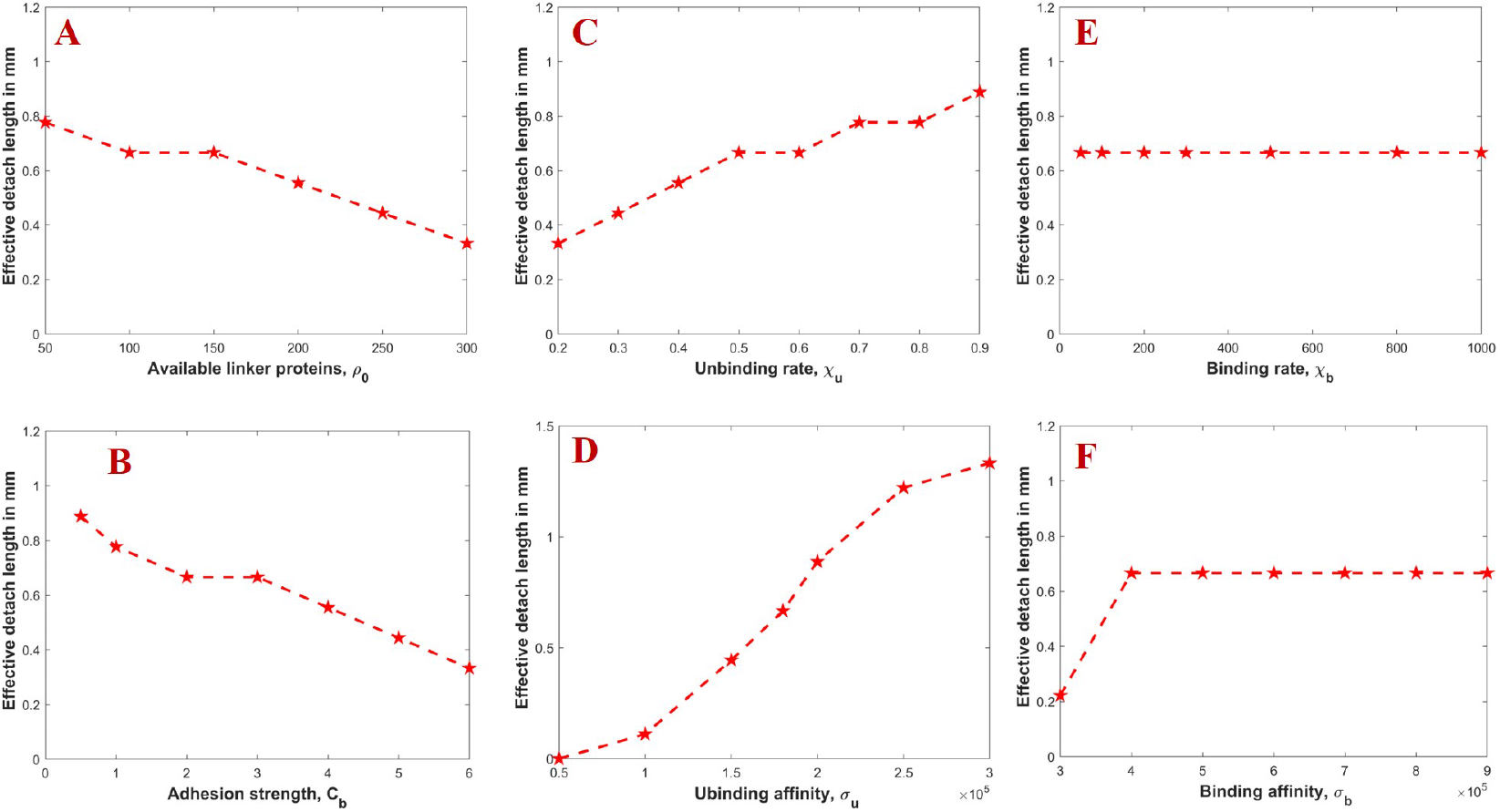
Molecular properties: (A) and (B) show the effects of adhesion bond density and adhesion bond strength on the effective detached length, respectively; (C) and (D) illustrate how increases in the unbinding rate and unbinding affinity affect the progression of detachment; (E) shows that the binding rate has little to no impact on the effective detached length; while (F) demonstrates that increasing the binding affinity reduces detachment progression.

The progression of detachment only occur when adhesion bonds binding the NL to the RPE are broken. The tendancy for these proteins to dissociate or unbind from their binding site determines how quickly the detachment will progress. Figure 13C and D show that increasing either the unbinding rate or the unbinding affinity of adhesion proteins accelerates detachment progression, leading to a greater effective detached length. In contrast, Figure 13E indicates that the binding rate has negligible influence on detachment, likely because reattachment (or healing) occurs only when the detached layers are in close proximity for a sufficient amount of time.

From a modeling perspective, the binding affinity is inversely related to the parameter *σ*_*b*_ in our formulation (Equation (13)); decreasing *σ*_*b*_ increases the binding affinity. As seen in Figure 13F, increasing the binding affinity reduces the rate of detachment progression. Despite the short simulation window, healing occurred when *σ*_*b*_ was sufficiently small, as further illustrated in Figure 14.

**Figure 14:**
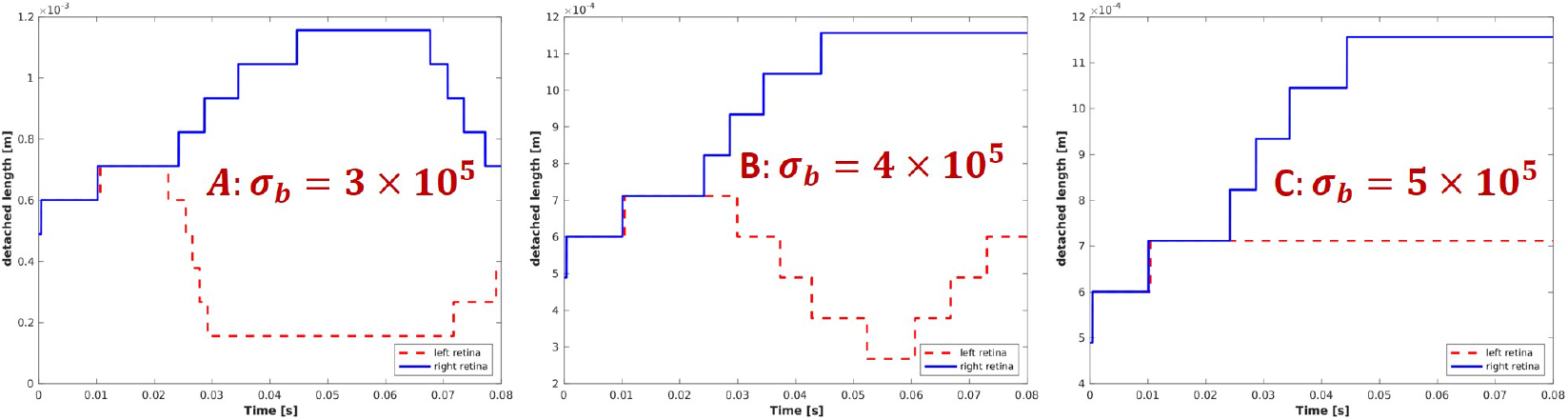
Detachment profile for binding affinity: The figure shows that decreasing *σ*_*b*_, which corresponds to increasing the binding affinity of adhesion proteins, can lead to healing even within a short simulation time.

From a quantitative perspective, these results suggest that enhanced binding affinity may contribute to biological reattachment. However, further experimental validation is necessary to confirm these findings in vivo.

#### 4.4.3. Goldstein Feedback Coefficients

Finally, we investigated the effect of the Goldstein feedback coefficients *α* and *β*, as defined in Equation (38), which determine the force the fluid exerts on the retina. Our results indicate that the spring coefficient *α* has a negligible effect on the progression of retinal detachment (RD). However, for the damping coefficient *β*, we observed that when |*β*| *>* 12, there is no noticeable effect on detachment progression. In contrast, for |*β*| *<* 12, the effective detached length increases as the magnitude of *β* increases. Since values for these parameters are well established in the literature—where *α* and *β* are typically chosen to be large and negative, such as (*α, β*) = (−100, −10) [52, 31], we conclude that under such conditions, these coefficients have minimal influence on the progression of detachment.

**Figure 15:**
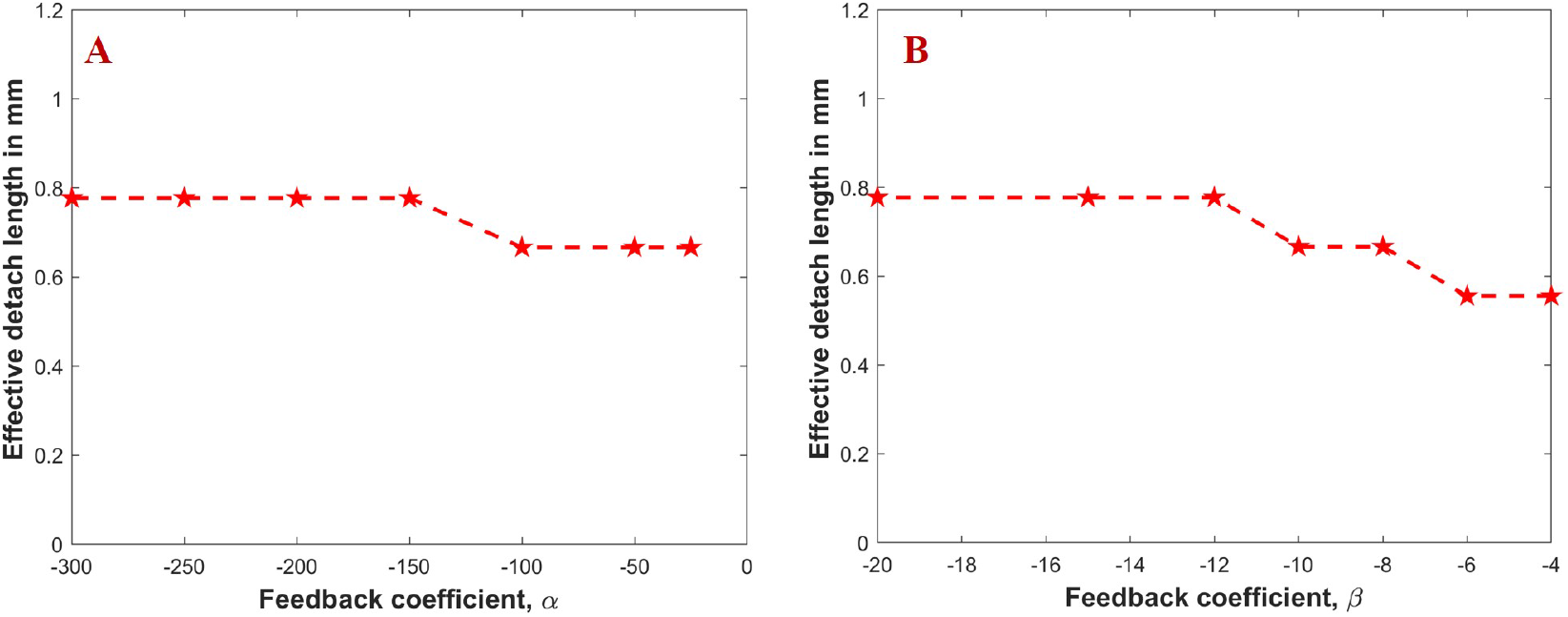
Goldstein feedback coefficients: The figure shows how the Goldstein feedback coefficients, which determine the force the fluid exerts on the retina, influence the progression of detachment. (A) and (B) respectively show the influence of *α* and *β* on the progression of detachment.

## 5. Conclusion

This work builds on the fluid-structure interaction (FSI) model, a modeling framework commonly used to study the interaction between fluid and deformable structures. In developing the retinal progression–fluid-structure interaction (RP-FSI) model to investigate the progression of retinal detachment, we incorporated into the FSI model a component that describes the adhesion bonds between the neural layer (NL) and the retinal pigmented epithelium (RPE). This component captures the dynamics of how these adhesion bonds are broken, resulting in further detachment.

We outlined the initial and boundary conditions of the model and provided a brief description of the numerical approach used for its solution. Since most existing fluid solvers were developed *ad hoc* for specific problems, integrating the adhesion dynamics model into them proved challenging. As a result, we developed a custom numerical algorithm in MATLAB to simulate the model. The MATLAB code used to simulate the model is publicly available on GitHub and can be accessed at: https://github.com/williamannan/Progression-of-retinal-detachment. Numerical results were then presented to illustrate how various model parameters influence the progression of retinal detachment.

Specifically, we found that increasing the eyewall velocity (*u*_*max*_), the initial detached length (*l*_0_), the elevation of the detached retina (*φ*), the retinal density (*ρ*_*s*_), the unbinding affinity of the adhesion proteins (*σ*_*u*_), and the unbinding rate (*χ*_*u*_) results in an increased effective detached length. Conversely, increasing the thickness of the retina (*h*^*^), the modulus of elasticity (*E*), the density of available adhesion proteins (*ρ*_0_), the fluid density (*ρ*_*f*_), and the adhesion bond strength (*C*_*b*_) reduces the effective detached length or slows down the progression. Finally, we observed that the dynamic fluid viscosity (*µ*), the binding rate (*χ*_*b*_), and the binding affinity (*σ*_*b*_) have minimal influence on the progression of retinal detachment.

While partial information on some model parameters exists in the literature, key values—such as adhesion bond strength, binding and unbinding affinities, and the distribution of available adhesion proteins—are not readily reported in forms directly applicable to our equations. To address this, we used available data as reference points and conducted simulations to examine how variations in these parameters affect model behavior. The baseline values used in this study, listed in Table 1, were selected based on a combination of available literature and simulation trends. However, future experimental studies are necessary to obtain more precise parameter estimates and further validate the model’s predictions.

The current model is formulated in two dimensions, with one axis along the retinal plane and the other perpendicular to the eyewall to represent fluid height. This 2D simplification limits the model’s ability to capture the full curvature and surface area of the retina; consequently, only horizontal eye rotations were considered. Notably, previous studies have shown that individuals with myopia are at higher risk of retinal detachment [21, 48]. Future work will focus including the curvature of the eyeball and extending the model to three dimensions to enhance physiological relevance.

## Supporting information

supplementary code for numerical code

## Acknowledgments

The authors would like to thank Prof. Joseph Skufca, Dr. James Greene, and Dr. Bethany Almeida for serving on William’s Ph.D. committee. Their valuable insights, constructive suggestions, and generous contributions greatly enhanced the quality of this work.

## Appendix A. Derivation of the Adhesion Protein Model

In this section, we derive an equation for the distribution of the adhesion proteins, denoted by *ρ*_*b*_(*s, t*), during retinal detachment. Let 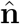 and 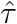 represent the unit normal and tangent vectors to the retina, respectively. Let **X**(*s, t*) denote any point along the retina and **X**^*e*^(*s*) the resting configuration or position of the retina before detachment. Define *ν* to be a unit vector perpendicular to the eyewall. The rate of change of the adhesion protein density *ρ*_*b*_(*s, t*) at any time is proportional to the difference between the binding and unbinding rates of the adhesion proteins. Hence, we express this as [5]:

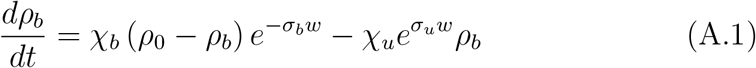

where *w* = | (**X** − **X**^*e*^) · *ν* | represents the vertical displacement of the retina, with *σ*_*b*_, *σ*_*u*_ *>* 0 describing the binding and unbinding affinities of the adhesion proteins respectively. Since the retina is represented as a two-dimensional structure, any point on the retina is given by **X**(*s, t*) = [*x*(*s, t*), *y*(*s, t*)]. Moreover, because the distribution of the adhesion proteins *ρ*_*b*_(*s, t*) depends on the position of the retina, the total derivative of *ρ*_*b*_ with respect to *t* is given by:

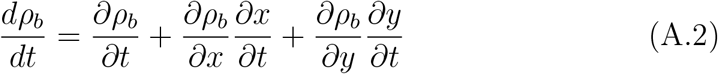

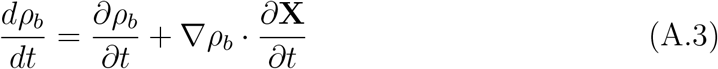

The expression in Equation (A.3) highlights that the rate of change of the adhesion protein density at any point along the retina depends not only on the biochemical binding and unbinding processes but also on how the retina’s motion affects the proteins distribution across space. The first term on the right-hand side, 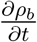 represents the local change in ρb at a fixed position, while the second term, 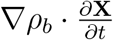, captures the advection effect, describing how the retina’s motion influences the spatial distribution of the adhesion proteins. Because the adhesion proteins bind the NL of the retina to the RPE, the distribution of *ρ*_*b*_ varies only along the retina and not across it. This implies that the gradient of *ρ*_*b*_ in the normal direction is zero, that is,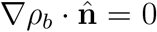.

Consequently, the gradient of *ρ*_*b*_ can be expressed as

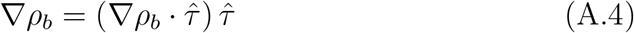

where 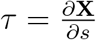 is a tangent vector along the retina. Substituting Equation (A.4) into Equation (A.3), we obtain

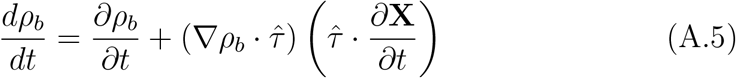

For any fixed time, the derivative of *ρ*_*b*_ with respect to the arc-length parameter *s* is given by

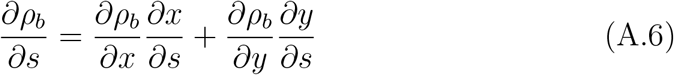

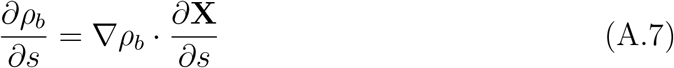

Substituting Equation (A.4) into Equation (A.7) yields

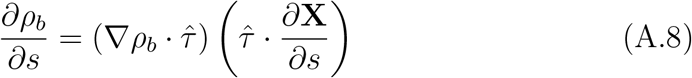

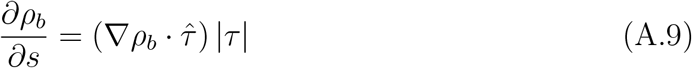

Thus, the gradient of the adhesion proteins is determined by only the tangential component and is given by

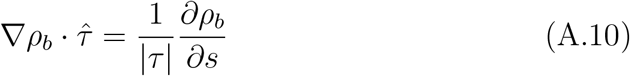

Substituting Equation (A.10) into Equation (A.5) gives

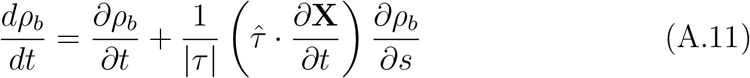

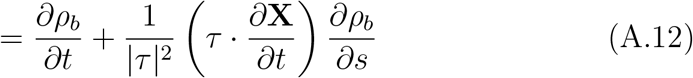

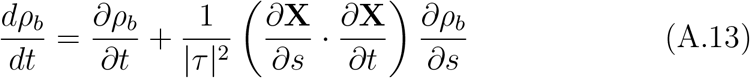

Substituting Equation (A.13) into Equation (A.1) gives

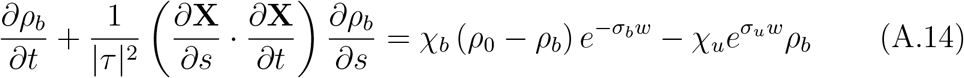

subject to the initial condition:

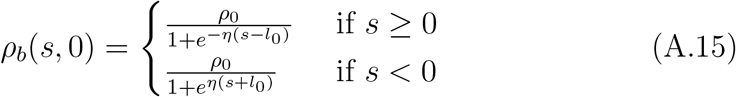

where *l*_0_ represent the initial detached length and *η*, some constant. Equation (A.14) is an advection reaction equation with the advection speed given by

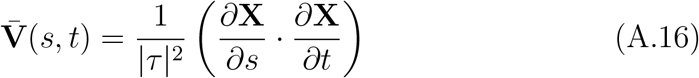

The model suggests that the change in the distribution of the adhesion proteins which determines the progression of the retinal detachment is advective in nature with the advection speed dependent on the geometry and the local speed of the retina.

