## supplementary code for numerical code for "Mathematical Model for the Progression of Rhegmatogenous Retinal Detachment (RRD)"

### 1. Introduction

In this supplementary material, we provide a detailed description of the numerical scheme used to solve the Retinal Progression–Fluid Structure Interaction (RP-FSI) model developed to study the progression of retinal detachment. The RP-FSI model consists of a coupled system of equations: the incompressible Navier–Stokes equations governing the fluid flow, equation describing the motion of the retina, and an equation modeling the distribution of adhesion bonds between the neural layer (NL) and the retinal pigmented epithelium (RPE).

To solve the Navier–Stokes equations, we used a non-iterative $P_2$ projection method. The retinal motion equation is solved using a finite difference scheme, while the adhesion bond distribution equation is handled using an implicit upwind method.

Figure 1 presents a flowchart of the overall numerical algorithm, illustrating how the scheme is advanced in time. In the following sections, we provide detailed explanations of each sub-component, including the discretization strategies and the solution procedures for each part of the model.

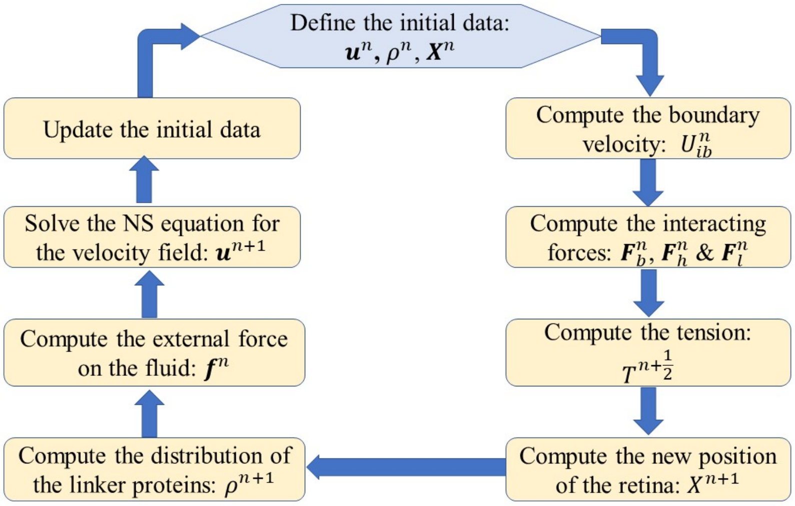


Figure 1: **Algorithm flowchart:** The figure shows time stepping algorithm flowchart used to solve the RP-FSI model

### 2. The Retinal Progression–Fluid Structure Interaction (RP-FSI) Model

Here, we present a summary of the nondimensionalized form of the RP-FSI model, together with the boundary conditions for the retinal position and tension. The incompressible Navier–Stokes equations describing the fluid flow are given by:

$$\frac{\partial \mathbf{u}}{\partial t} + (\mathbf{u} \cdot \nabla)\,\mathbf{u} + \nabla p - \frac{1}{Re}\nabla^2 \mathbf{u} = \mathbf{f}(\mathbf{x}, t) \tag{1}$$

$$\nabla \cdot \mathbf{u} = 0 \tag{2}$$

where $Re = \frac{\rho_f u_{max} L_0}{\mu}$ is the Reynolds number, and the external force the retina exert on the fluid given by

$$\mathbf{f}(\mathbf{x}, t) = \rho \int_\Gamma \mathbf{F}_l(s,t)\delta\left(\mathbf{x} - \mathbf{X}\right) ds, \tag{3}$$

with $\rho = \frac{\rho_d}{\rho_f L_0}$.

The motion of the retina in response to the fluid flow is governed by:

$$\frac{\partial^2 \mathbf{X}}{\partial t^2} = \frac{\partial}{\partial s}\left(T\frac{\partial \mathbf{X}}{\partial s}\right) + \mathbf{F}_b(s,t) - \mathbf{F}_l(s,t) - \mathbf{F}_h(s,t) \tag{4}$$

The tension $T$ generated within the retina satisfies the equations:

$$\frac{\partial \mathbf{X}}{\partial s}\cdot\frac{\partial^2}{\partial s^2}\left(T\frac{\partial \mathbf{X}}{\partial s}\right)=\frac{1}{2}\frac{\partial^2}{\partial t^2}\left(\frac{\partial \mathbf{X}}{\partial s}\cdot\frac{\partial \mathbf{X}}{\partial s}\right)-\frac{\partial^2 \mathbf{X}}{\partial s\partial t}\cdot\frac{\partial^2 \mathbf{X}}{\partial s\partial t}-\frac{\partial \mathbf{X}}{\partial s}\cdot\frac{\partial}{\partial s}\left(\mathbf{F}_b-\mathbf{F}_l-\mathbf{F}_h\right) \tag{5}$$

and

$$\frac{\partial \mathbf{X}}{\partial s}\cdot\frac{\partial \mathbf{X}}{\partial s}=1 \tag{6}$$

The interacting forces are given by:

$$\mathbf{F}_b(s,t)=-\frac{\partial^2}{\partial s^2}\left(\gamma\frac{\partial^2 \mathbf{X}}{\partial s^2}\right) \tag{7}$$

$$\mathbf{F}_h(s,t)=C_b\rho_b(s,t)\left[(\mathbf{X}-\mathbf{X}^e)\cdot\nu\right]\nu \tag{8}$$

$$\mathbf{F}_l(s,t)=\alpha\int_0^t\left(U_{ib}(s,t)-\frac{\partial \mathbf{X}}{\partial \tau}\right)d\tau+\beta\left(U_{ib}(s,t)-\frac{\partial \mathbf{X}}{\partial t}\right) \tag{9}$$

where $\gamma$ is the dimensionless bending resistance, $\mathbf{X}^e$ is the resting configuration of the retina (i.e., the position of the retina before detachment), $\nu$ is a unit vector perpendicular to the eyewall, $C_b$ is the strength of adhesion bonds, and

$$\mathbf{U}_{ib}(s,t)=\int_\Omega \mathbf{u}(\mathbf{x},t)\delta_h(\mathbf{x}-\mathbf{X})d\mathbf{x}, \tag{10}$$

is the boundary fluid velocity at the fluid–structure interface. Here, $\delta(\mathbf{x})$ is a two-dimensional Dirac delta function defined as in [5]:

$$\delta_h(\mathbf{x})=\frac{1}{h^2}\phi\left(\frac{x_1}{h}\right)\phi\left(\frac{x_2}{h}\right), \tag{11}$$

where

$$\phi(r)=\begin{cases}\frac{1}{8}\left(5-2|r|-\sqrt{-7+12|r|-4r^2}\right), & 1\le|r|\le 2,\\ \frac{1}{8}\left(3-2|r|+\sqrt{1+4|r|-4r^2}\right), & 0\le|r|<1,\\ 0, & \text{otherwise.}\end{cases} \tag{12}$$

The equation governing the density distribution of adhesion proteins binding the NL to the RPE is given by:

$$\frac{\partial \rho_b}{\partial t}+\frac{1}{|\tau|^2}\left(\frac{\partial \mathbf{X}}{\partial s}\cdot\frac{\partial \mathbf{X}}{\partial t}\right)\frac{\partial \rho_b}{\partial s}=\chi_b\left(\rho_0-\rho_b\right)e^{-\sigma_b w}-\chi_u e^{\sigma_u w}\rho_b \tag{13}$$

where $\rho_0$ is the density of available adhesion proteins (i.e., the density of adhesion proteins before detachment), and $\chi_b$ and $\chi_u$ are the binding and unbinding rates, respectively. The parameters $\sigma_b$ and $\sigma_u$ denote the binding and unbinding affinities of the adhesion proteins.

The retinal position and the tension generated within the retina are subject to the following boundary conditions. At the detached end of the retina, we assume both the curvature and the shear stress are zero, leading to the boundary condition [4]:

$$\left.\frac{\partial^2 \mathbf{X}}{\partial s^2}\right|_{s=0} = \mathbf{0}, \tag{14}$$

$$\left.\frac{\partial^3 \mathbf{X}}{\partial s^3}\right|_{s=0} = \mathbf{0} \tag{15}$$

At the far end, we assume the retina remains attached to the eyewall such that the tangent vector is horizontal. Moreover, within the short simulation time, the entire retina within the domain cannot completely detach. This gives the boundary condition:

$$\left.\frac{\partial \mathbf{X}}{\partial s}\right|_{s=\pm L} = (\pm 1,\ 0), \tag{16}$$

$$\left.\mathbf{X}\right|_{\pm L} = \mathbf{X}_0 \tag{17}$$

In 3D, both halves of the detached retina are connected. To mimic this, we impose the boundary condition:

$$\left.T\right|_{s=0} = \pm \kappa_{\Sigma} |(\mathbf{X}_r - \mathbf{X}_l)| \tag{18}$$

on the tension at the detached end, where $\kappa_{\Sigma}$ is a spring constant that depends on the stiffness of the retina, and $\mathbf{X}_r$ and $\mathbf{X}_l$ represent the right and left halves of the detached retina, respectively.

At the extreme ends, since the retina remains bound to the eyewall, we assume its acceleration matches that of the eyewall. Applying this condition to Equation (4) results in:

$$\left.\frac{\partial}{\partial s}\left(T \frac{\partial \mathbf{X}}{\partial s}\right)\right|_{s=\pm L} = A_p + (\mathbf{F}_l + \mathbf{F}_h - \mathbf{F}_b) \tag{19}$$

where $A_p$ denotes the acceleration of the eyewall.

### 3. Numerical Solution to RP-FSI model

Detailed numerical scheme and discretization process used to solve the RP-FSI model is presented here. Figure 1 provides a schematic overview of the main computational steps involved in the time-stepping solution of the model.

To solve the fluid equations (i.e., *the Navier–Stokes equations*), we employ the non-iterative $P_2$ projection method. Spatial derivatives are approximated using finite differences on a staggered grid. The Navier–Stokes solver is advanced in time using an explicit Adams–Bashforth scheme for the nonlinear advection term, while the viscous diffusion term is treated implicitly using the Crank–Nicolson method. This results in an implicit–explicit Crank–Nicolson (IMEX-CN) time-stepping scheme.

The adhesion protein density equation, as presented in Equation (13), is discretized using an implicit upwind scheme to ensure stability in the advection-dominated regime.

The retinal deformation equation given in Equation (4) is solved using a finite difference method, treating the first term on the right-hand side implicitly and the remaining terms explicitly.

The overall strategy is to use the known solution at the current time step to compute the solution at the next time step.

#### *3.1. Discretization of the Boundary Velocity and the Interacting Forces*

Let $\mathbf{x} \in \mathbb{R}^2$ be an Eulerian coordinate in the fluid domain and $\mathbf{X} \in \mathbb{R}^2$, a Lagrangian coordinate describing the position on the retina. The fluid velocity at the boundary of the retina given by Equation (10) is approximated using numerical quadrature at time $t_n$ as

$$U_{ib}^n = \sum_{(x_i,x_j)\in\mathbf{x}} \mathbf{u}^n \delta_h(\mathbf{x}-\mathbf{X}^n)h^2, \tag{20}$$

where $h$ is the spatial step size, and the summation is taken over the entire fluid domain. For any fixed point in the fluid domain $\mathbf{x} = (x, y)$ and any fixed point on the retina $\mathbf{X} = (X_1, X_2)$, the discrete Dirac delta function is given by

$$\delta_h(\mathbf{x}-\mathbf{X}^n) = \frac{1}{h^2}\phi\left(\frac{x - X_1^n}{h}\right)\phi\left(\frac{y - X_2^n}{h}\right). \tag{21}$$

Assuming the bending rigidity of the retina, denoted by $\gamma$, is constant, the bending force given by Equation (7) can be discretized using central difference approximations and applying the retinal boundary conditions given by Equation (14) to (17)

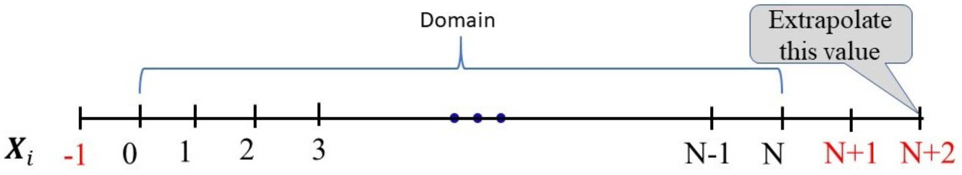


Figure 2: **Partition for the retinal domain**: The figure shows the partition of the retinal domain used to solve for the position of the retina as it interacts with the fluid.

Considering one half of the retina (e.g., the right half), we obtain from Equation (14):

$$\frac{\mathbf{X}_1 - 2\mathbf{X}_0 + \mathbf{X}_{-1}}{(\triangle s)^2} = \mathbf{0} \implies \mathbf{X}_{-1} = 2\mathbf{X}_0 - \mathbf{X}_1. \tag{22}$$

Similarly, from equation (16), we get:

$$\mathbf{X}_{N+1} = \mathbf{X}_{N-1} + 2\triangle s(\pm 1, 0). \tag{23}$$

We then extrapolate the value of $\mathbf{X}$ at $i = N + 2$. The discretized form of the bending force becomes:

$$\mathbf{F}_b\Big|_i = -\gamma \frac{\mathbf{X}_{i+2} - 4\mathbf{X}_{i+1} + 6\mathbf{X}_i - 4\mathbf{X}_{i-1} + \mathbf{X}_{i-2}}{(\triangle s)^4}, \quad i = 1, 2, \cdots, N. \tag{24}$$

For $i = 0$, we apply the boundary condition given in equation (15), which yields:

$$\frac{D_{ss}\mathbf{X}\Big|_1 - D_{ss}\mathbf{X}\Big|_{-1}}{2\triangle s} = \mathbf{0} \implies D_{ss}\mathbf{X}\Big|_1 = D_{ss}\mathbf{X}\Big|_{-1}. \tag{25}$$

Thus, the bending force at $i = 0$ becomes:

$$\mathbf{F}_b\Big|_0 = -\gamma \frac{D_{ss}\mathbf{X}_1 - 2D_{ss}\mathbf{X}_0 + D_{ss}\mathbf{X}_{-1}}{(\triangle s)^2}. \tag{26}$$

But since $D_{ss}\mathbf{X}_0 = \mathbf{0}$ and $D_{ss}\mathbf{X}_{-1} = D_{ss}\mathbf{X}_1$, we finally have:

$$\mathbf{F}_b\Big|_0 = -2\gamma \frac{\mathbf{X}_1 - 2\mathbf{X}_0 + \mathbf{X}_{-1}}{(\triangle s)^4}. \tag{27}$$

The force $\mathbf{F}_l(s,t)$ exerted by the fluid on the retina, given by the Goldstein feedback law [3],

$$\mathbf{F}_l(s,t) = \alpha \int_0^t \left( U_{ib}(s,t') - \frac{\partial \mathbf{X}}{\partial t'} \right) dt' + \beta \left( U_{ib}(s,t) - \frac{\partial \mathbf{X}}{\partial t} \right),$$

is discretized as

$$\mathbf{F}_l^n = \alpha \sum_{k=1}^{n} \left( U_{ib}^k - \frac{\mathbf{X}^k - \mathbf{X}^{k-1}}{\triangle t} \right) \triangle t + \beta \left( U_{ib}^n - \frac{\mathbf{X}^n - \mathbf{X}^{n-1}}{\triangle t} \right). \tag{28}$$

Having defined the density of the adhesion proteins $\rho_b^n$ at time $t_n = t_0 + n\triangle t$, the adhesion force restraining the motion of the retina can be expressed in the form

$$\mathbf{F}_h^n = C_b \rho_b^n \left[ (\mathbf{X}^n - \mathbf{X}^e) \cdot \nu \right] \nu, \tag{29}$$

where $\mathbf{X}^e$ is the resting position of the retina (i.e., the position of the retina before detachment), and $\nu$ is a unit vector perpendicular to the eyewall.

#### *3.2. Discretization of the Tension Equation*

The tension $T$, generated within the retina as the fluid pushes against it satisfies Equation (5) and also subject to the inextensibility constraint:

$$\frac{\partial \mathbf{X}}{\partial s} \cdot \frac{\partial \mathbf{X}}{\partial s} = 1.$$

Theoretically, the first term on the right-hand side of Equation (5) vanishes. However, numerical errors arising from enforcing the inextensibility constraint during the computation necessitate its retention to maintain numerical stability [6]. Instead of discarding this term, we ensure that the updated position of the retina at the next time step, $\mathbf{X}^{n+1}$, satisfies the inextensibility constraint. In the scheme, the tension force is computed at intermediate time steps, denoted as $T^{n+\frac{1}{2}}$, at intermediate nodes $i + \frac{1}{2}$, as illustrated in Figure 3.

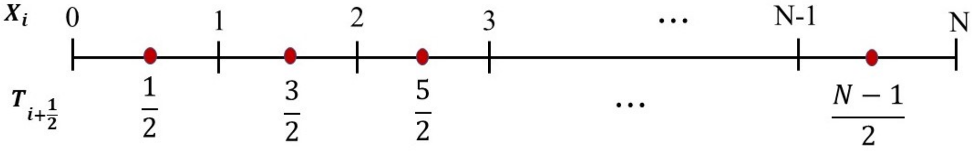


Figure 3: **Partition for the Tension**: The figure shows the partition for the retinal domain used to solve to the tension in the retina as it interact with the fluid

The goal is to express the tension equation in the matrix form

$$AT^{n+\frac{1}{2}} = \mathbf{B}, \tag{30}$$

where $A$ and $\mathbf{B}$ are $(N-1)\times(N-1)$ matrix and $N-1$ vector obtained from the left and the right hand side of the tension equation respectively.

Let

$$Y_{i+\frac{1}{2}} = D_s\mathbf{X}|_{i+\frac{1}{2}} = \frac{\mathbf{X}_{i+1} - \mathbf{X}_i}{\triangle s}, \tag{31}$$

then the left hand side of Equation (5) can be express in a discrete form as

$$\begin{aligned} D_s\mathbf{X}\cdot D_{ss}\left(TDs\mathbf{X}\right)_{i+\frac{1}{2}} &= Y_{i+\frac{1}{2}} D_{ss}\left(TY\right)_{i+\frac{1}{2}} \\ &= Y_{i+\frac{1}{2}} \cdot \frac{T_{i+\frac{3}{2}}Y_{i+\frac{3}{2}} - 2T_{i+\frac{1}{2}}Y_{i+\frac{1}{2}} + T_{i-\frac{1}{2}}Y_{i-\frac{1}{2}}}{(\triangle s)^2} \\ &= \frac{T_{i+\frac{3}{2}}Y_{i+\frac{1}{2}}\cdot Y_{i+\frac{3}{2}} - 2T_{i+\frac{1}{2}}|Y_{i+\frac{1}{2}}|^2 + T_{i-\frac{1}{2}}Y_{i+\frac{1}{2}}\cdot Y_{i-\frac{1}{2}}}{(\triangle s)^2} \end{aligned}$$

Thus, for $i = 1, 2, \ldots, N-2$,

$$D_s\mathbf{X}\cdot D_{ss}\left(TDs\mathbf{X}\right)_{i+\frac{1}{2}} = \frac{1}{(\triangle s)^2}\left[T_{i+\frac{3}{2}}Y_{i+\frac{1}{2}}\cdot Y_{i+\frac{3}{2}} - 2T_{i+\frac{1}{2}}|Y_{i+\frac{1}{2}}|^2 + T_{i-\frac{1}{2}}Y_{i+\frac{1}{2}}\cdot Y_{i-\frac{1}{2}}\right]. \tag{32}$$

At $i = 0$, we apply centered difference approximation on the first interval giving

$$D_s\mathbf{X}\cdot D_{ss}\left(TDs\mathbf{X}\right)_{\frac{1}{2}} = Y_{\frac{1}{2}}\cdot\frac{D_s(TY)_1 - D_s(TY)_0}{(\triangle s)} \tag{33}$$

$$= Y_{\frac{1}{2}}\cdot\frac{T_{\frac{3}{2}}Y_{\frac{3}{2}} - T_{\frac{1}{2}}Y_{\frac{1}{2}}}{(\triangle s)^2} - Y_{\frac{1}{2}}\cdot\frac{T_{\frac{1}{2}}Y_{\frac{1}{2}} - T_0Y_0}{\frac{1}{2}(\triangle s)^2} \tag{34}$$

which simplifies to

$$D_s\mathbf{X}\cdot D_{ss}\left(TDs\mathbf{X}\right)_{\frac{1}{2}} = \frac{1}{(\triangle s)^2}\left[T_{\frac{3}{2}}Y_{\frac{3}{2}}\cdot Y_{\frac{1}{2}} - 3T_{\frac{1}{2}}|Y_{\frac{1}{2}}|^2\right] + 2T_0Y_{\frac{1}{2}}\cdot Y_0 \tag{35}$$

with the second term on the right accounting for the boundary condition for the tension at the detached end of the retina when RRD develop from retinal hole where $T_0$ is given by Equation (18). In the case of great retinal tear, $T_0 = 0$.

Similarly, at $i = N-1$,

$$D_s\mathbf{X}\cdot D_{ss}(TD_s\mathbf{X})\Big|_{N-\frac{1}{2}} = Y_{N-\frac{1}{2}}\cdot\frac{D_s(TD_s\mathbf{X})_N - D_s(TD_s\mathbf{X})_{N-1}}{(\triangle s)}$$
$$= \frac{Y_{N-\frac{1}{2}}}{(\triangle s)}\cdot D_s(TD_s\mathbf{X})_N - \left[\frac{T_{N-\frac{1}{2}}|Y_{N-\frac{1}{2}}|^2 - T_{N-\frac{3}{2}}Y_{N-\frac{1}{2}}\cdot Y_{N-\frac{3}{2}}}{(\triangle s)^2}\right]$$

which simplifies to

$$D_s\mathbf{X}\cdot D_{ss}(TD_s\mathbf{X})\Big|_{N-\frac{1}{2}} = -\frac{1}{(\triangle s)^2}\left[T_{N-\frac{1}{2}}|Y_{N-\frac{1}{2}}|^2 - T_{N-\frac{3}{2}}Y_{N-\frac{1}{2}}\cdot Y_{N-\frac{3}{2}}\right] + \frac{Y_{N-\frac{1}{2}}}{(\triangle s)}\cdot D_s(TD_s\mathbf{X})_N \tag{36}$$

But form Equation (19),

$$D_s(TD_s\mathbf{X})\Big|_N = A_p - (\mathbf{F}_b - \mathbf{F}_l - \mathbf{F}_h)\Big|_{s=\pm L} \tag{37}$$

where $A_p$ is the acceleration of the eyewall. From Equation (32), Equation (35), and Equation (36), the left hand side of the tension equation equation can be written in the form

$$AT^{n+\frac{1}{2}} + \mathbf{F}_{BC} \tag{38}$$

where

$$\mathbf{F}_{BC} = 2T_0Y_{\frac{1}{2}}\cdot Y_0 + \frac{Y_{N-\frac{1}{2}}}{(\triangle s)}\cdot D_s(TD_s\mathbf{X})_N \tag{39}$$

represent boundary terms that are incoperated into the right hand vector $\mathbf{B}$ of Equation (30).

The first term on the right-hand side of the tension equation is discretized as

$$\frac{1}{2}D_{tt}\left(D_s\mathbf{X}\cdot D_s\mathbf{X}\right) = \frac{\left(D_s\mathbf{X}\cdot D_s\mathbf{X}\right)^{n+1} - 2\left(D_s\mathbf{X}\cdot D_s\mathbf{X}\right)^n + \left(D_s\mathbf{X}\cdot D_s\mathbf{X}\right)^{n-1}}{2(\triangle t)^2} \tag{40}$$

Enforcing the inextensibility constraint on the position of the retina at future time $t_{n+1} = t_0 + (n+1)\triangle t$, that is

$$\left(D_s\mathbf{X}\cdot D_s\mathbf{X}\right)^{n+1} = 1. \tag{41}$$

gives

$$\frac{1}{2}D_{tt}\left(D_s\mathbf{X}\cdot D_s\mathbf{X}\right) = \frac{1}{2(\triangle t)^2}\left[1 - 2\left(D_s\mathbf{X}\cdot D_s\mathbf{X}\right)^n + \left(D_s\mathbf{X}\cdot D_s\mathbf{X}\right)^{n-1}\right], \tag{42}$$

The right hand side vector of Equation (30) becomes

$$\mathbf{B} = \frac{1}{2} D_{tt} \left( D_s \mathbf{X} \cdot D_s \mathbf{X} \right) - D_{st} \mathbf{X} \cdot D_{st} \mathbf{X} - D_s \mathbf{X} \cdot D_s \left( \mathbf{F}_b - \mathbf{F}_l - \mathbf{F}_h \right) - \mathbf{F}_{BC}. \quad (43)$$

Thus, the tension at the next half time step is given by

$$T^{n+\frac{1}{2}} = A^{-1} \mathbf{B}. \quad (44)$$

where the entries of the matrix $A$ are given by $\mathbf{Y}_{i+\frac{1}{2}} \cdot \mathbf{Y}_{j+\frac{1}{2}}$ for $i, j = 0, 1, 2, 3, \ldots, N-1$

*3.3. Discretization for Retinal/Material Equation*

The nondimensionalized form of the model describing the motion of the retina in responds to the fluid flow is given by Equation (4). In Section 3.1, we detailed how to discretize the forces $\mathbf{F}_b^n, \mathbf{F}_l^n$, and $\mathbf{F}_h^n$. The general approach to solving this equation is to rewrite it in the form

$$\frac{\mathbf{X}^{n+1} - 2\mathbf{X}^n + \mathbf{X}^{n-1}}{(\triangle t)^2} = M \mathbf{X}^{n+1} + \bar{\mathbf{F}} \quad (45)$$

where $M$ is an $N \times N$ matrix derived from the first term on the right-hand side of Equation (4). We now detail how to construct the matrix $M$. At this stage, we assume that the tension $T^{n+\frac{1}{2}}$ has already been computed, as outlined in Section 3.2, since the entries of $M$ depends on $T$.

Consider one-half of the detached retina (e.g., the right half), partitioned into $N$ equal segments as illustrated in Figure 2. The tension term in the retinal equation can then be discretized as

$$D_s(T D_s \mathbf{X}) \Big|_i = \frac{T_{i+\frac{1}{2}} D_s \mathbf{X} \Big|_{i+\frac{1}{2}} - T_{i-\frac{1}{2}} D_s \mathbf{X} \Big|_{i-\frac{1}{2}}}{(\triangle s)} \quad (46)$$

$$= \frac{T_{i+\frac{1}{2}} \left( \mathbf{X}_{i+1} - \mathbf{X}_i \right) - T_{i-\frac{1}{2}} \left( \mathbf{X}_i - \mathbf{X}_{i-1} \right)}{(\triangle s)^2}. \quad (47)$$

Thus, for $i = 1, 2, \ldots, N-1$, we obtain

$$D_s(T D_s \mathbf{X}) \Big|_i = \frac{1}{(\triangle s)^2} \left[ T_{i+\frac{1}{2}} \mathbf{X}_{i+1} - \left( T_{i+\frac{1}{2}} + T_{i-\frac{1}{2}} \right) \mathbf{X}_i + T_{i-\frac{1}{2}} \mathbf{X}_{i-1} \right]. \quad (48)$$

At $i = 0$, we employ a forward difference approximation:

$$D_s(TD_s\mathbf{X})\Big|_0 = \frac{T_{\frac{1}{2}} D_s \mathbf{X}_{\frac{1}{2}} - T_0 D_s \mathbf{X}_0}{\frac{1}{2}(\triangle s)} \tag{49}$$

$$= \frac{2T_{\frac{1}{2}}(\mathbf{X}_1 - \mathbf{X}_0)}{(\triangle s)^2} - \frac{2T_0 D_s \mathbf{X}|_0}{(\triangle s)}. \tag{50}$$

From Equation (22), we know that $\mathbf{X}_{-1} = 2\mathbf{X}_0 - \mathbf{X}_1$. Therefore, using a centered difference approximation for $D_s\mathbf{X}_0$, we obtain

$$D_s\mathbf{X}_0 = \frac{\mathbf{X}_1 - \mathbf{X}_{-1}}{2(\triangle s)} = \frac{\mathbf{X}_1 - \mathbf{X}_0}{(\triangle s)}. \tag{51}$$

Substituting Equation (51) into Equation (50) yields

$$D_s(TD_s\mathbf{X})\Big|_0 = \frac{2T_{\frac{1}{2}}\mathbf{X}_1 - 2T_{\frac{1}{2}}\mathbf{X}_0}{(\triangle s)^2} - \frac{2T_0(\mathbf{X}_1 - \mathbf{X}_0)}{(\triangle s)^2}, \tag{52}$$

where $T_0$ is the boundary tension condition at the detached end given by Equation (18). The second term on the right-hand side of Equation (52) is absorbed into the force term $\bar{\mathbf{F}}$ in Equation (45). At the far end, we assume the retina remains attached to the eyewall, such that

$$D_{tt}\mathbf{X}\Big|_{s=\pm L} = A_p, \tag{53}$$

where $A_p$ represents the acceleration of the eyewall. This leads to

$$D_s(TD_s\mathbf{X})\Big|_{i=N} = A_p - (\mathbf{F}_b - \mathbf{F}_l - \mathbf{F}_h)\Big|_N. \tag{54}$$

Since $D_s(TD_s\mathbf{X})|_{i=N}$ does not explicitly depend on $\mathbf{X}$, Equation (54) is also incorporated into $\bar{\mathbf{F}}$ in Equation (45). As a result, the final row of matrix $M$ consists entirely of zeros, making $M$ a singular matrix. With the matrix $M$ defined, Equation (45) can now be rewritten as

$$\left[I - (\triangle t)^2 M\right] \mathbf{X}^{n+1} = 2\mathbf{X}^n - \mathbf{X}^{n-1} + (\triangle t)^2 \bar{\mathbf{F}}, \tag{55}$$

where

$$\bar{\mathbf{F}} = \mathbf{F}_b^n - \mathbf{F}_l^n - \mathbf{F}_h^n - \frac{2T_0(\mathbf{X}_1 - \mathbf{X}_0)}{(\triangle s)^2} + D_s(TD_s\mathbf{X})\Big|_{i=N}. \tag{56}$$

The updated position of the retina is then given by

$$\mathbf{X}^{n+1} = \left[I - (\triangle t)^2 M\right]^{-1} \left(2\mathbf{X}^n - \mathbf{X}^{n-1} + (\triangle t)^2 \bar{\mathbf{F}}\right). \tag{57}$$

*3.4. Discretization of Adhesion Protein Equation*

The density distribution of the adhesion proteins is governed by Equation (13). To discretize this equation, we apply an implicit upwind scheme under the assumption that the advection speed is positive. While this assumption may not always hold, a modified scheme would be required when the direction of advection changes. Referring to Figure 2, and recalling that $\mathbf{X}_{-1} = 2\mathbf{X}_0 - \mathbf{X}_1$, the advection speed at the $i^{\text{th}}$ node is expressed as:

$$\bar{V}_i = \frac{1}{|\tau|^2} \left( D_s\mathbf{X} \cdot D_t\mathbf{X} \right) \Big|_i = \frac{D_s\mathbf{X}|_i \cdot D_t\mathbf{X}|_i}{D_s\mathbf{X}|_i \cdot D_s\mathbf{X}|_i} \tag{58}$$

where the temporal and spatial derivatives are given, respectively, by

$$D_t\mathbf{X}|_i = \frac{\mathbf{X}_i^n - \mathbf{X}_i^{n-1}}{\triangle t}, \tag{59}$$

and

$$D_s\mathbf{X}\Big|_i = \frac{\mathbf{X}_{i+1}^n - \mathbf{X}_{i-1}^n}{2\triangle s}, \quad \text{for } i = 0, 1, 2, \ldots, N-1. \tag{60}$$

At $i = N$, the boundary condition is defined as:

$$D_s\mathbf{X}\Big|_N = (\pm 1, 0), \tag{61}$$

corresponding to the right and left halves of the detached retina, respectively. With $\bar{V}_i$ now discretized, the advection equation describing the distribution of the adhesion proteins becomes:

$$\frac{\rho_{b_i}^{n+1} - \rho_{b_i}^n}{\triangle t} + \bar{V}_i \frac{\rho_{b_i}^{n+1} - \rho_{b_{i-1}}^{n+1}}{\triangle s} = \chi_b \left( \rho_0 - \rho_{b_i}^{n+1} \right) e^{-\sigma_b w} - \chi_u e^{\sigma_u w} \rho_{b_i}^{n+1} \tag{62}$$

Multiplying through by $\triangle t$ and grouping terms yields:

$$\xi_1 \rho_{b_i}^{n+1} + \xi_2 \rho_{b_{i-1}}^{n+1} = \triangle t \chi_b \rho_0 e^{-\sigma_b w} + \rho_{b_i}^n \tag{63}$$

where the coefficients $\xi_1$ and $\xi_2$ are defined as:

$$\xi_1 = 1 + \frac{\triangle t}{\triangle s} \bar{V}_i + \triangle t \left[ \chi_b e^{-\sigma_b w} + \chi_u e^{\sigma_u w} \right] \tag{64}$$

$$\xi_2 = -\frac{\triangle t}{\triangle s} \bar{V}_i \tag{65}$$

Equation (63) can then be expressed in matrix form:

$$[A_\rho] \rho_b^{n+1} = B_\rho \tag{66}$$

which can be solved to obtain the updated density distribution of the adhesion proteins.

#### 3.5. Solving the Navier-Stokes Equations

The motion of the fluid is governed by the incompressible Navier–Stokes equations, given in Equation (1) and (2). Various numerical schemes have been developed to solve these equations. In this work, we adopt a non-iterative $P_2$ projection method [1, 2]. To advance the solution in time, the advection term is treated explicitly using the Adams–Bashforth method, while the diffusion term is handled implicitly using the Crank–Nicolson method. Spatial discretization is performed using finite differences on a staggered grid.

We begin by discretizing the external force exerted by the retina on the fluid, and then present the time-stepping procedure for solving the Navier–Stokes equations. The spatial discretization, which is also based on finite differences on a staggered grid, is discussed in detail afterward.

The external force exerted on the fluid by the retina is given by

$$\mathbf{f}(\mathbf{x}, t) = \rho \int_{\Gamma} \mathbf{F}_l(s, t)\delta\left(\mathbf{x} - \mathbf{X}(s, t)\right)\, ds.$$

Its discrete approximation is given by:

$$\mathbf{f}^n_{i,j} = \rho \sum_{k=1}^{N} \mathbf{F}^n_{l_k} \delta_h\left(\mathbf{x}_{i,j} - \mathbf{X}^n_k\right) h, \tag{67}$$

where $\delta_h(\mathbf{x} - \mathbf{X}^n_k)$ is the discrete Dirac delta function, as defined in Equation (21).

##### 3.5.1. Time stepping procedure for $P_2$ projection method

Before describing the numerical solution procedure for the $P_2$ projection method, Let's briefly review the concept of operator splitting, which forms the foundation of the time-stepping scheme used. Suppose we want to solve the equation

$$\frac{\partial w}{\partial t} = g_1(x, t) + g_2(x, t), \tag{68}$$

subject to the initial condition $w(x, 0) = w_0$. In operator splitting, the right-hand side is decomposed into two parts, which are solved sequentially over each time step. Introducing an intermediate solution $w^*$, we discretize the equation as follows:

$$\frac{w^* - w^n}{\Delta t} = g_1(x, t), \tag{69}$$

$$\frac{w^{n+1} - w^*}{\Delta t} = g_2(x, t). \tag{70}$$

Here, $w^n$ represents the solution at the $n$-th time step. First, Equation (69) is solved using $w^n$ to obtain the intermediate solution $w^*$; this intermediate value is then used as the initial condition to solve Equation (70) for $w^{n+1}$. This process is repeated recursively over time.

The spatial derivatives in the momentum equations (ie *advection and diffusion terms*) are approximated using finite difference schemes on a staggered grid. Details of the scheme, as well as the grid arrangement and the discretization procedure will be discussed in the following subsections.

A summary of the $P_2$ projection scheme is presented in the flowchart in Figure 4. Being a multi-step method, it requires initial velocities at two time levels. The initial velocity $\mathbf{u}^0$ is provided, and the velocity $\mathbf{u}^1$ is obtained by integrating Equation (1) explicitly over the first time interval $[t_0, t_0+\Delta t]$ using a very small time step $dt_f = \frac{\Delta t}{N_t}$, where $N_t$ is the number of sub-iterations. The resulting velocity is taken as $\mathbf{u}^1$, and the associated pressure is denoted by $p^{n-\frac{1}{2}}$.

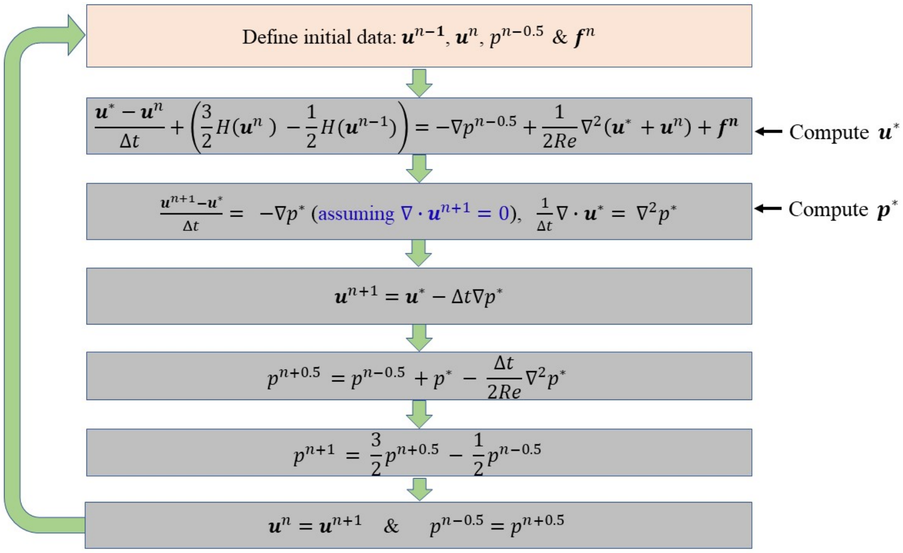


Figure 4: **Flowchart for a $P_2$ projection method**: The figure shows a summary of how a $P_2$ projection method is applied in solving the Navier–Stokes equations.

Given $\mathbf{u}^{n-1}$, $\mathbf{u}^n$, and $p^{n-\frac{1}{2}}$, the right-hand side of the momentum equation of the Navier–Stokes equations, given by Equation (1), is decomposed by applying operator splitting. The advection term is treated using the explicit Adams–Bashforth method, while the diffusion term is treated implicitly using the Crank–Nicolson scheme. Assuming that the pressure in the momentum equation can be expressed as

$$p = p^{n-0.5} + \tilde{p}, \tag{71}$$

where $\tilde{p}$ is an unknown pressure correction that can be used to estimate the true pressure at the next time step. Then, introducing an intermediate velocity field $\mathbf{u}^*$ and applying operator splitting, the discretization of the momentum equation becomes:

$$\frac{\mathbf{u}^* - \mathbf{u}^n}{\Delta t} + \left(\frac{3}{2}H(\mathbf{u}^n) - \frac{1}{2}H(\mathbf{u}^{n-1})\right) = -\nabla p^{n-0.5} + \frac{1}{2Re}\nabla^2\left(\mathbf{u}^* + \mathbf{u}^n\right) + \mathbf{f}^n, \tag{72}$$

and

$$\frac{\mathbf{u}^{n+1} - \mathbf{u}^*}{\Delta t} = -\nabla\tilde{p}, \tag{73}$$

where $H(\mathbf{u}) = (\mathbf{u}\cdot\nabla)\mathbf{u}$ denotes the advection term. Here, $\mathbf{u}^*$ may not satisfy the continuity condition given by Equation (2). Discretization of $H(\mathbf{u})$ is done using finite difference appromiximation on a staggered grid to be discussed in detail in the next sub-section.

Equation (72) can be written in the form:

$$\mathbf{u}^* - \frac{\Delta t}{2Re}\nabla^2\mathbf{u}^* = \text{RHS}, \tag{74}$$

where

$$\text{RHS} = \mathbf{u}^n - \Delta t\left(\frac{3}{2}H(\mathbf{u}^n) - \frac{1}{2}H(\mathbf{u}^{n-1})\right) - \Delta t\nabla p^{n-0.5} + \frac{\Delta t}{2Re}\nabla^2\mathbf{u}^n + \Delta t\mathbf{f}^n. \tag{75}$$

Since all the terms on the right-hand side are known, Equation (74) can be solved for $\mathbf{u}^*$.

**Note**: In two dimensional (*respectively 3D*) flow, Equation (74) must be decomposed as a system of equations given by

$$u^* - \frac{\Delta t}{2Re}\left(\frac{\partial^2 u^*}{\partial x^2} + \frac{\partial^2 u^*}{\partial y^2}\right) = \text{RHS}_u \tag{76}$$

$$v^* - \frac{\Delta t}{2Re}\left(\frac{\partial^2 v^*}{\partial x^2} + \frac{\partial^2 v^*}{\partial y^2}\right) = \text{RHS}_v \tag{77}$$

where each component of the velosity field is solved separately.

Assuming that the velocity at the next time step $\mathbf{u}^{n+1}$ satisfies the continuity condition, i.e.,

$$\nabla \cdot \mathbf{u}^{n+1} = 0, \tag{78}$$

taking the divergence of Equation (73) and applying Equation (78), we can derive the following Poisson equation for the pressure correction:

$$\nabla^2 \tilde{p} = \frac{1}{\Delta t} \nabla \cdot \mathbf{u}^*, \tag{79}$$

and solve for $\tilde{p}$. Once $\tilde{p}$ is obtained, the velocity field at the next time level can be computed from Equation (73) as:

$$\mathbf{u}^{n+1} = \mathbf{u}^* - \Delta t \nabla \tilde{p}. \tag{80}$$

Having solved for $\tilde{p}$, $\mathbf{u}^*$, and $\mathbf{u}^{n+1}$, we derive an expression for the pressure at the next half time step (i.e., $p^{n+0.5}$). To achieve this, we re-write Equation (73) as:

$$\mathbf{u}^* = \mathbf{u}^{n+1} + \Delta t \nabla \tilde{p}. \tag{81}$$

Substituting Equation (81) into Equation (72) and simplifying, we recover:

$$\frac{\mathbf{u}^{n+1} - \mathbf{u}^n}{\Delta t} + \left( \frac{3}{2} H(\mathbf{u}^n) - \frac{1}{2} H(\mathbf{u}^{n-1}) \right) = -\nabla p^{n+\frac{1}{2}} + \frac{1}{2Re} \nabla^2 \left( \mathbf{u}^{n+1} + \mathbf{u}^n \right) + \mathbf{f}^n, \tag{82}$$

where $p^{n+0.5}$ is defined as:

$$p^{n+\frac{1}{2}} = p^{n-0.5} + \tilde{p} - \frac{\Delta t}{2Re} \nabla^2 \tilde{p}. \tag{83}$$

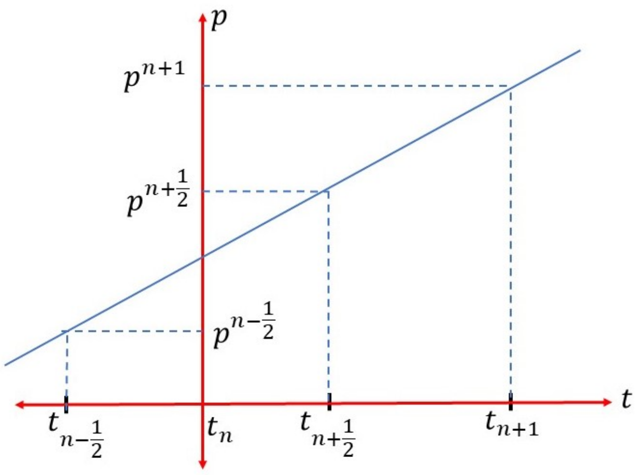


Figure 5: **Pressure extrapolation**: The figure shows a simple sketch used to extrapolate the fluid pressure at the next time step

Finally, using the midpoint pressures $p^{n-\frac{1}{2}}$ and $p^{n+\frac{1}{2}}$, the pressure at the next time step is extrapolated as:

$$p^{n+1} = \frac{3}{2}p^{n+\frac{1}{2}} - \frac{1}{2}p^{n-\frac{1}{2}}. \tag{84}$$

*3.5.2. Spatial Discretization*

Here, we discuss how the spatial discretization can be carried out. We start with the discretization of the advection term before the diffusion term. Given the velocity field $\mathbf{u} = (u, v)$, the nonlinear advection term in two dimensions can be expressed as:

$$(\mathbf{u} \cdot \nabla)\, \mathbf{u} = \left( u\frac{\partial u}{\partial x} + v\frac{\partial u}{\partial y}, \quad u\frac{\partial v}{\partial x} + v\frac{\partial v}{\partial y} \right) \tag{85}$$

The spatial derivatives are approximated using a finite difference scheme on a staggered grid, as shown in Figure 6. Here, the $u$ and $v$ components of the velocity field are defined at the horizontal and vertical edges of the computational cell respectively, while the pressure is defined at the center of the cell.

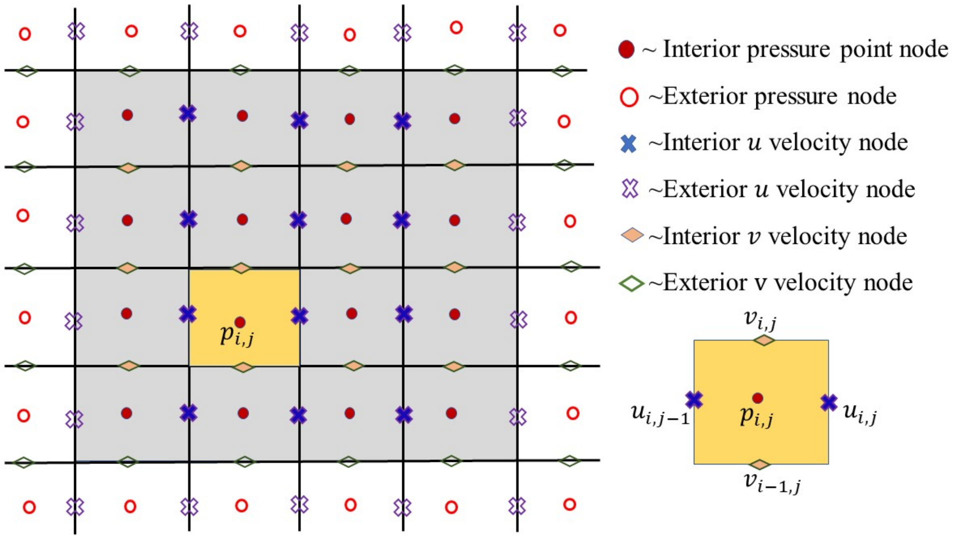


Figure 6: **Staggered grid**: The figure shows a staggered grid used to approximate the spatial derivatives.

The general approach for discretizing this term involves the following steps:

- Compute the partial derivatives at the edges of the **velocity cells** for each component of the velocity.

- Average these edge-based derivatives to estimate the derivative at the center of each velocity cell.

It is important to distinguish between **computational cells** and **velocity cells**. The velocity cells are centered at the edges of the computational cells, as illustrated in Figure 7, and this staggered arrangement helps enhance numerical stability [7].

**Discretization for the $u$-component:** We begin with the discretization of the term $u\frac{\partial u}{\partial x}$. This term is computed at the center of the $u$-velocity cell, which lies at a vertical edge of the computational cell. To compute it, we first evaluate:

$$uu_x\Big|_{i,j+\frac{1}{2}} = \frac{u_{i,j+1} + u_{i,j}}{2} \cdot \frac{u_{i,j+1} - u_{i,j}}{\triangle x} \tag{86}$$

$$uu_x\Big|_{i,j-\frac{1}{2}} = \frac{u_{i,j} + u_{i,j-1}}{2} \cdot \frac{u_{i,j} - u_{i,j-1}}{\triangle x} \tag{87}$$

Then, the value of $u\frac{\partial u}{\partial x}$ at the center of the $u$-velocity cell is approximated by averaging the above two expressions:

$$uu_x\Big|_{i,j} = \frac{uu_x|_{i,j+\frac{1}{2}} + uu_x|_{i,j-\frac{1}{2}}}{2} \tag{88}$$

Next, we discretize the term $v\frac{\partial u}{\partial y}$. This term involves computing the rate of change of $u$ in the vertical direction using vertical edges of the computational cell. The intermediate values are given by:

$$vu_y\Big|_{i+\frac{1}{2},j} = \frac{v_{i,j+1} + v_{i,j}}{2} \cdot \frac{u_{i+1,j} - u_{i,j}}{\triangle y} \tag{89}$$

$$vu_y\Big|_{i-\frac{1}{2},j} = \frac{v_{i-1,j+1} + v_{i-1,j}}{2} \cdot \frac{u_{i,j} - u_{i-1,j}}{\triangle y} \tag{90}$$

Averaging these expressions gives the discrete approximation for $v\frac{\partial u}{\partial y}$ at the center of the $u$-velocity cell:

$$vu_y\Big|_{i,j} = \frac{vu_y|_{i+\frac{1}{2},j} + vu_y|_{i-\frac{1}{2},j}}{2} \tag{91}$$

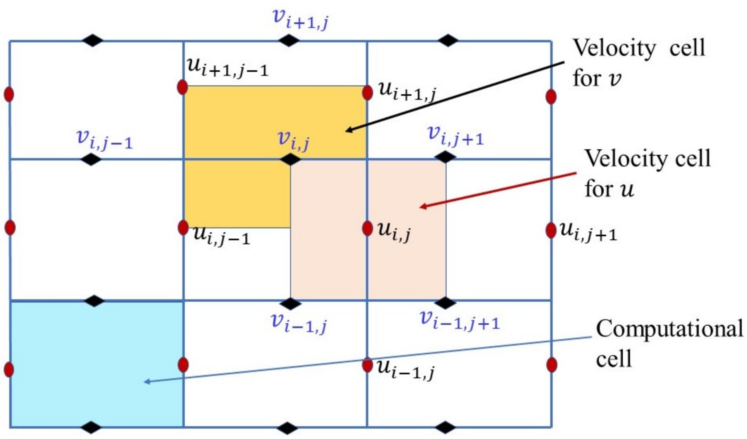


Figure 7: **Computation and velocity cells**: The figure a staggered grid representation of computational and velocity cells.

**Discretization for the $v$-component:** The discretization of the advection term for the $v$-component of the velocity field is similar and follows the same procedure. We begin with the term $u\frac{\partial v}{\partial x}$, which involves computing horizontal derivatives of $v$. At the edges of the $v$-velocity cell:

$$uv_x\Big|_{i,j+\frac{1}{2}} = \frac{u_{i+1,j} + u_{i,j}}{2} \cdot \frac{v_{i,j+1} - v_{i,j}}{\triangle x} \tag{92}$$

$$uv_x\Big|_{i,j-\frac{1}{2}} = \frac{u_{i+1,j-1} + u_{i,j-1}}{2} \cdot \frac{v_{i,j} - v_{i,j-1}}{\triangle x} \tag{93}$$

Then, at the center of the $v$-velocity cell:

$$uv_x\Big|_{i,j} = \frac{uv_x|_{i,j+\frac{1}{2}} + uv_x|_{i,j-\frac{1}{2}}}{2} \tag{94}$$

The final term is $v\frac{\partial v}{\partial y}$, which is computed at the horizontal edges of the $v$-velocity cell:

$$vv_y\Big|_{i+\frac{1}{2},j} = \frac{v_{i+1,j} + v_{i,j}}{2} \cdot \frac{v_{i+1,j} - v_{i,j}}{\triangle y} \tag{95}$$

$$vv_y\Big|_{i-\frac{1}{2},j} = \frac{v_{i,j} + v_{i-1,j}}{2} \cdot \frac{v_{i,j} - v_{i-1,j}}{\triangle y} \tag{96}$$

And the average gives the approximation:

$$vv_y\Big|_{i,j} = \frac{vv_y|_{i+\frac{1}{2},j} + vv_y|_{i-\frac{1}{2},j}}{2} \tag{97}$$

*3.5.3. Discretization of the Laplacian*

The Laplacian operator is discretized using a finite difference scheme as follows:

$$\nabla^2\psi = \frac{\psi_{i,j-1} - 2\psi_{i,j} + \psi_{i,j+1}}{(\triangle x)^2} + \frac{\psi_{i-1,j} - 2\psi_{i,j} + \psi_{i+1,j}}{(\triangle y)^2} \tag{98}$$

With this definition, when solving either the $u$ or $v$ component of the Navier Stokes equations, or the Poisson equation for the pressure correction $\tilde{p}$ given by Equation (79), you simply replace the function $\psi$ with the corresponding function $u$, $v$, or $\tilde{p}$ respectively.

In practical implementation, all these expressions should be computed in a vectorized form to enhance computational efficiency. The staggered grid layout helps reduce numerical oscillations and is widely used in immersed boundary methods and fluid solvers. The MATLAB code developed to solve the model is publicly available on GitHub and can be accessed at: `https://github.com/williamannan/Progression-of-retinal-detachment`

### Acknowledgments

The authors would like to thank Prof. Joseph Skufca, Dr. James Greene, and Dr. Bethany Almeida for serving on William's Ph.D. committee. Their valuable insights, constructive suggestions, and generous contributions greatly enhanced the quality of this work.

### Declaration of interest: none
